# RNAs undergo phase transitions with lower critical solution temperatures

**DOI:** 10.1101/2022.10.17.512593

**Authors:** Gable M. Wadsworth, Walter J. Zahurancik, Xiangze Zeng, Paul Pullara, Lien B. Lai, Vaishnavi Sidharthan, Rohit V. Pappu, Venkat Gopalan, Priya R. Banerjee

**Author notes:** These authors contributed equally (Gable M. Wadsworth, Walter J. Zahurancik, Xiangze Zeng). Correspondence should be addressed to: P.R.B; R.V.P; V.G.

## Abstract

Co-phase separation of RNAs and RNA-binding proteins is thought to drive the biogenesis of ribonucleoprotein granules. RNAs can also undergo phase transitions in the absence of proteins. However, the physicochemical driving forces of protein-free, RNA-driven phase transitions remain unclear. Here, we report that RNAs of various types undergo phase transitions with system-specific lower critical solution temperatures (LCSTs). This entropically-driven phase behavior requires Mg^2+^ ions and is an intrinsic feature of the phosphate backbone that is modulated by RNA bases. RNA-only condensates can additionally undergo enthalpically favorable percolation transitions within dense phases. This is enabled by a combination of Mg^2+^-dependent bridging interactions among phosphate groups and RNA base-stacking / base-pairing. Phase separation coupled to percolation can cause dynamical arrest of RNAs within condensates and can suppress the catalytic activity of an RNase P ribozyme. Our work highlights the need to incorporate RNA-driven phase transitions into models for RNP granule biogenesis.

## Introduction

Membraneless biomolecular condensates are macromolecular compartments that form via spontaneous or driven phase transitions aided by homotypic and heterotypic interactions involving multivalent protein and RNA molecules ^1, 2, 3, 4, 5, 6, 7, 8^. The driving forces that govern the formation of biomolecular condensates have largely been investigated through protein engineering-aided alterations of protein-protein and protein-RNA interactions ^9, 10, 11, 12, 13, 14, 15^. However, key studies on phase transitions of RNA-binding proteins (RBPs) have highlighted the critical role of RNAs ^16, 17, 18, 19^. For example, RBPs undergo phase separation based on RNA-to-protein ratios. At a low RNA-to-protein stoichiometry, RNAs facilitate RBP phase separation, whereas the same RBPs form soluble ribonucleoprotein (RNP) complexes at high RNA-to-protein ratios ^20^. Importantly, RNAs have also been reported to undergo phase separation in the absence of proteins ^21, 22, 23, 24, 25^. However, the role of homotypic RNA-RNA interactions that drive protein-free RNA phase transitions is less well understood than the determinants of phase transitions in RNA-protein mixtures driven by heterotypic interactions ^26, 27, 28, 29, 30^.

RNA condensation *in vitro*, defined either as the compaction of single RNA molecules or phase separation at higher concentrations, depends on polyvalent cations (e.g., spermine, poly-lysine, cobalt (III) hexamine, Mg^2+^ ions), crowding agents (e.g., polyethylene glycol), and/or RNA denaturation ^22, 23, 25, 31, 32, 33^. Notably, in the presence of divalent cations such as Mg^2+^, GC-rich RNAs with trinucleotide repeats undergo a sol-gel transition, *i.e*., percolation *in vitro*. This is thought to underlie the formation of intracellular RNA granules in RNA repeat expansion disorders ^25, 34, 35, 36^. Percolation transitions of RNA molecules refer to the formation of system-spanning networks driven by the physical crosslinking of cohesive motifs known as stickers ^37, 38^. In the case of RNA molecules, the stickers are the nucleobases, and the physical crosslinks involve a combination of base pairing and base stacking interactions ^24^. Percolation on its own can give rise to the formation of thermoreversible, physical gels ^39^. Percolation, which is a networking transition defined by associative interactions, can be coupled to phase separation, which is a segregative transition driven mainly by effective polymer-solvent interactions ^38, 40, 41, 42, 43, 44, 45, 46^. Since RNA condensates represent the formation of dilute and dense coexisting phases, phase separation coupled to percolation (PSCP) is likely to underlie RNA condensate formation.

The current working model for the formation of RNA-based condensates is that the relevant phase transitions are driven by base-pairing interactions ^31, 47^. The contributions of base pairs should weaken with increasing temperature. This is because of the Boltzmann weighting of base pair energies as a function of temperature. If true, then a direct consequence of the current working model is that RNA-RNA interactions must be suppressed at high temperatures ^48, 49, 50, 51, 52^, and condensate formation by RNAs must feature system-specific upper critical solution temperatures (UCSTs) ^42, 53, 54^. In this scenario, for an RNA concentration that places the system in the two-phase regime at one temperature, an increase in temperature beyond its apparent UCST will cause condensate dissolution and exit from the two-phase regime. In contrast, if an increase in temperature transitions the system from a one-phase regime to a two-phase regime, then the phase transition is defined by a lower critical solution temperature (LCST). Such transitions are entropically driven, become favorable with increasing temperature, and are primarily mediated by the release of solvent molecules, which are mixtures of water and solution ions.

Are phase transitions of RNA molecules enthalpically driven UCST-type processes, entropically driven LCST-type processes, or a combination of the two? To answer this question, we performed heating and cooling temperature ramps on a series of RNAs of varying sequence and size and monitored changes in mesoscale morphology with a light microscope. Our investigations revealed that in the presence of divalent salts, RNA phase separation is entropically driven (favored as temperature increases) with the occurrence of system specific LCSTs in the presence of Mg^2+^ ions. The LCST-type transitions of RNAs are enabled by the phosphate backbone, driven by desolvation entropy, and tuned by the RNA-RNA and RNA-solvent interactions in a nucleobase-specific fashion. Within the dense phases that form via LCST-type phase separation, the RNA molecules can undergo percolation transitions. These appear to be driven by enthalpically favorable intermolecular interactions including base pairing and stacking that give rise to droplet-spanning networks. Percolation within dense phases can lead to dynamical arrest of RNA condensates. This is tied to the strengths of physical crosslinks, which determine the timescales for making and breaking of crosslinks ^55, 56^, and the interplay of these dynamics with molecular transport. RNA systems featuring LCST-type phase separation and percolation within dense phases, that also undergo dynamical arrest, will not always fully transition back to the homogeneous single-phase regime even when cooled well below their apparent LCSTs ^24, 56^. This can give rise to hysteretic phase behavior.

By titrating solution conditions as well as RNA sequence complexity and composition, we have identified determinants of entropically driven phase separation that is coupled to enthalpically driven percolation. Moreover, through experiments on a ribozyme (the catalytic RNA subunit of RNase P), we demonstrate that PSCP has an inhibitory effect on RNA catalysis. We ascribe this outcome to the percolation-induced dynamical arrest that engenders entanglement of ribozymes within condensates. Our findings of a broadly applicable LCST-type transition in different RNAs suggest a molecular basis for the biogenesis of various stress-responsive ribonucleoprotein (RNP) condensates. They also highlight the need to incorporate intrinsic RNA phase transitions into emerging models for the formation and regulation of RNP condensates.

## Results

### Homopolymeric RNAs undergo temperature dependent phase transitions

As early as 1967, poly(rA) RNA molecules were observed to undergo condensation in a narrow range of temperatures becoming compact at higher temperatures ^21^. More recently, other nucleic acid homopolymers have been reported to undergo phase separation in various conditions, typically with multivalent cations and/or crowding agents ^23, 57, 58^. We recently reported that poly(rU) displays thermoresponsive phase behavior with UCST- and LCST-type transitions in the presence of multivalent cations such as spermine ^22^. While the UCST transition can be conceptualized through the formation of multivalent ion-induced physical crosslinking of RNA chains, the origin of an LCST transition is more nuanced and points to a delicate interplay between RNA-RNA and solvent-mediated interactions ^59^ that remain poorly understood. Here, we used temperature-controlled microscopy to study different RNA polymers and test the hypothesis that LCST-type phase separation behavior might extend to other RNAs (**Fig. 1a-d**).

**Figure 1:**
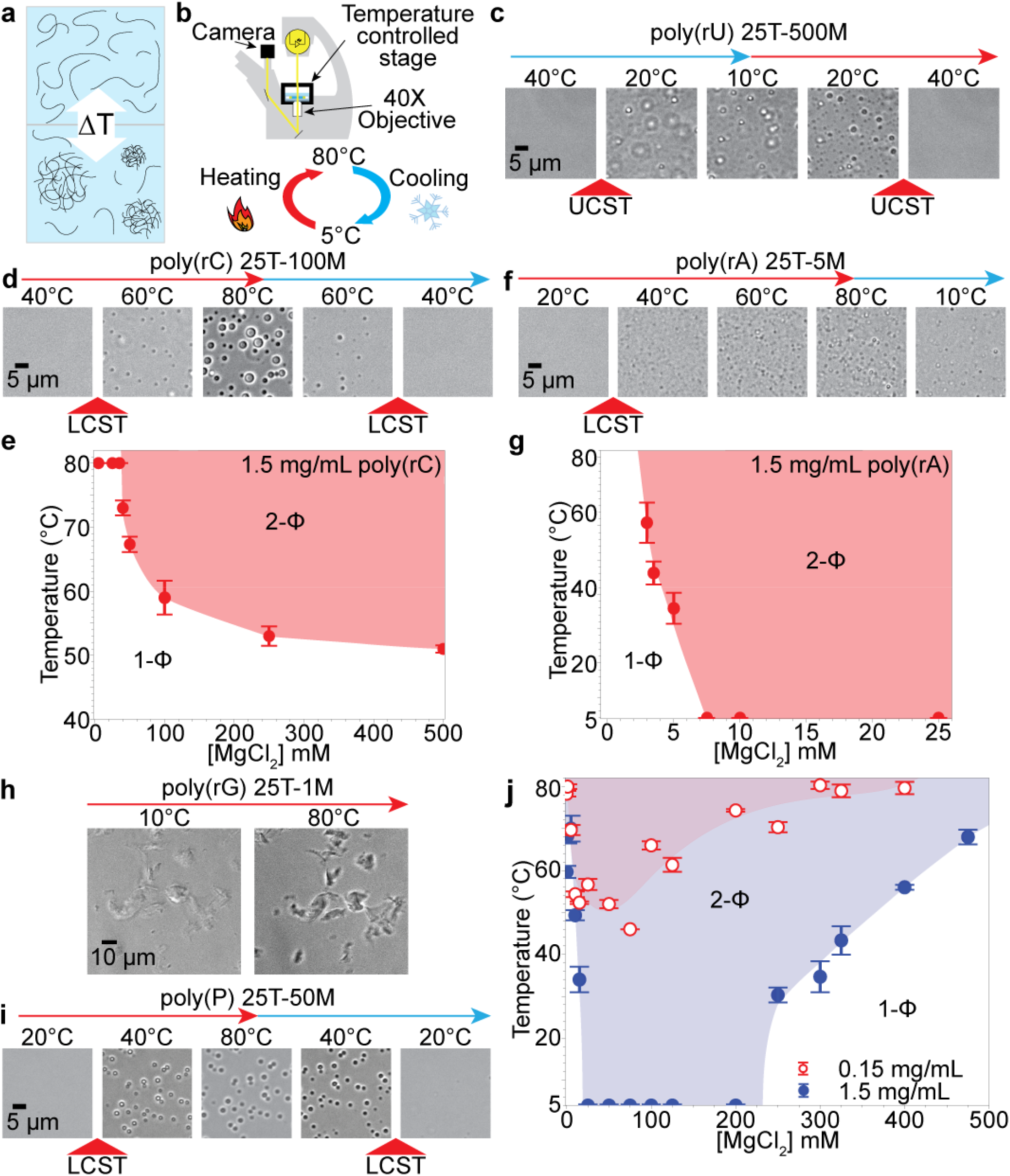
Temperature-controlled phase separation of RNA homopolymers. **(a)** Depiction of temperature-dependent RNA phase separation. **(b)** A schematic of temperature-controlled microscopy assay to probe RNA phase separation. **(c)** Phase separation of poly(rU) samples upon cooling suggesting a UCST-type phase behavior. Brightfield images of 1.5 mg/mL poly(rU) in 25 mM Tris-HCl (pH 7.5 at 25°C), 500 mM Mg^2+^ (25T-500M buffer) during heating (red arrow) and cooling (cyan arrow) as indicated; upper cloud-point temperature (UCPT) of this sample is 25.2 ± 1.2°C. The corresponding state diagram of poly(rU) is shown in Supplementary Fig. 4. **(d)** Phase separation of poly(rC) samples upon heating suggesting an LCST-type phase behavior. Brightfield images of 1.5 mg/mL poly(rC) in 25 mM Tris-HCl (pH 7.5 at 25°C), 100 mM Mg^2+^ (25T-100M buffer) during heating (red arrow) and cooling (cyan arrow) as indicated; lower cloud-point temperature (LCPT) of this sample is 57.6 ± 1.9°C. **(e)** State diagram of poly(rC) based on a series of experiments akin to those reported in panel **d**. **(f)** Phase separation and arrest of poly(rA) upon heating. Brightfield images of 1.5 mg/mL poly(rA) in 25 mM Tris-HCl (pH 7.5 at 25°C), 5 mM Mg^2+^ (25T-5M buffer) during heating (red arrow) and cooling (cyan arrow) as indicated; observed LCPT of this sample is 39.3 ± 8.5°C. Upon cooling, poly(rA) droplets did not dissolve. **(g)** State diagram of poly(rA) based on a series of experiments akin to those reported in panel **f**. **(h)** Brightfield images of 1.5 mg/mL poly(rG) in 25 mM Tris-HCl (pH 7.5 at 25°C), 1 mM Mg^2+^ (25T-1M buffer) at two different temperatures. Extensive irreversible aggregation is evident. **(i)** LCST-type phase separation of poly(P). Brightfield images of 1.5 mg/mL poly(P) in 25 mM Tris-HCl (pH 7.5 at 25°C), 250 mM Mg^2+^ (25T-250M buffer) during heating (red arrow) and cooling (cyan arrow) as indicated; observed LCPT of this sample is 35.3 ± 5.0°C. **(j)** State diagram of poly(P) based on a series of experiments akin to those reported in panel **i** using either 1.5 mg/mL (blue) and 0.15 mg/mL (red) poly(P) and a varying amount of Mg^2+^. Buffer notation used: the number in front of “T” indicates [Tris-HCl] and the number in front of “M” indicates [Mg^2+^] in each buffer.

We found a reversible UCST-type phase separation for poly(rU) in presence of Mg^2+^ ions^22, 60^ (**Fig. 1c**; **Supplementary Fig. S1**; **Movie 1**). However, under similar conditions, poly(rC) undergoes a reversible LCST-type phase transition with droplet formation at 57.6 ± 1.9°C (defined as T_phase_). The condensates dissolve when cooled to temperatures that are 1-2°C below T_phase_ (**Fig. 1d**; **Movie 2**). Poly(rA) also underwent an LCST-type phase transition with T_phase_ being 39.3 ± 8.5°C. However, the transition occurred at 1 mM Mg^2+^ instead of 100 mM Mg^2+^ required for phase separation of poly(rC). Importantly, poly(rA) condensates did not demonstrate reversibility and instead formed arrested condensates that persisted upon cooling below T_phase_ (**Fig. 1e**; **Movies 3, 4**). Arrest of poly(rA) condensates did not depend on the quench depth (*i.e*., the difference between the experimental temperature and the lower cloud point temperature of the sample) but was immediate upon crossing into the two-phase regime (**Supplementary Fig. S2**). Finally, in the case of poly(rG), we found significant irreversible mesoscale network formation under all conditions above 0.15 mM Mg^2+^ (**Fig. 1F**; **Supplementary Fig. S3**; **Movie 5**). This observation is consistent with a previous report on poly(rG) RNA ^23^. Overall, our data suggest that poly(rC) RNA can form condensates via a reversible LCST-type phase transition upon heating in the presence of Mg^2+^, whereas poly(rA) and poly(rG) RNAs form irreversible arrested phases. Both observations are concordant with a PSCP process ^61^, with the timescales for rearrangement of physical crosslinks being refractory to reversibility for poly(rA) condensates ^24^. Arrested networks are not the same as equilibrium solids ^24, 56^. Instead, they are metastable states derive from a dynamical interplay between long lived crosslinks that in turn suppress molecular transport ^55, 56^.

### Inorganic polyphosphate undergoes an LCST–type phase transition

To parse the determinants of RNA LCST phase behavior, we studied the phase behavior of inorganic polyphosphate, poly(P), a linear polyanion. We observed that poly(P) underwent phase separation at room temperature in the presence of Mg^2+^. Remarkably, we observed that 1.5 mg/mL poly(P) forms condensates with a reversible LCST-type transition **(Fig. 1i**; **Movie 6**) in a [Mg^2+^]-dependent manner. Away from the critical temperature, the observation of LCST behavior is discernible by the presence of a lower cloud point temperature (LCPT). We observed a LCPT for poly(P) systems and the precise value of LCPT changes non-monotonically as a function of Mg^2+^ concentration **(Fig. 1j**). These data suggest that the phase behavior of poly(P) with Mg^2+^ ions is reentrant, and phase separation is favored within a specific range of Mg^2+^ ion concentrations. Phase separation was observed across the entire range of accessible temperatures (5°C < T < 80°C) in our experiments when the [Mg^2+^] was between 25 mM and 250 mM with a fixed poly(P) concentration of 1.5 mg/mL. These observations suggest that the LCPT of these samples must be below 5°C. By lowering the concentration of the poly(P) to 0.15 mg/mL, we observed that the LCPT of poly(P) samples increased with reversible phase separation being observed at all Mg^2+^ ion concentrations tested **(Fig. 1j**). These data suggest that the negatively charged phosphate backbone of RNA can contribute to the observed LCST-type transition of RNA molecules in presence of Mg^2+^ ions.

### Entropic effects drive temperature-dependent, Mg^2+^ ion-induced chain compaction of polyphosphate

The occurrence of LCST-type phase separation is typical for polymers with hydrophobic moieties where the increased entropic penalty of solvation at high temperatures drives their decreased solubility with increased temperature. To understand the molecular driving forces of the Mg^2+^-dependent LCST-type transition of poly(P), we employed all-atom molecular dynamics simulations. For phase transitions driven by homotypic interactions, there is typically a strong coupling between the driving forces for single-chain coil-to-globule transitions and the driving forces for phase transitions ^59, 62, 63^. For polymers showing UCST phase behavior, a single chain in a solvent will undergo a globule-to-coil transition as the temperature increases ^62^. In this scenario, chains are compact at low temperature and adopt expanded conformations above a system-specific transition temperature (T_θ_ ≈ T_phase_). For systems with LCST phase behavior, the single chain prefers expanded conformations at lower temperatures, and undergoes chain compaction as temperature increases above T_θ_ ≈ T_phase_^62, 63^. We obtained a molecular picture of the impact of temperature on polyphosphate conformations by performing all-atom molecular dynamics simulations. The system comprises explicit representations of a polyphosphate molecule PO_3_^2-^-(PO_3_^-^)_16_-O-PO_3_^2^, water molecules, neutralizing Mg^2+^ ions, and excess salt ions whilst ensuring overall electroneutrality of the system. Details of the simulation setup are described in the *Materials and Methods* section and in the *Supporting Information* **(Supplementary Figs. S4-S6**; **Supplementary Table S2**).

The simulation results show that polyphosphate in 100 mM MgCl_2_ is more collapsed at 368 K than at 278 K **(Fig. 2a**). This is quantified in terms of the distributions of radii of gyration (R_g_), which show a clear shift toward compact conformations at the higher temperature. This observation is consistent with features of systems that show LCST phase behavior ^62, 63^. Next, we quantified the distribution of Mg^2+^ ions around polyphosphate atoms. We found that Mg^2+^ ions prefer to bind to the O2 (non-bridging) atoms instead of the OS (bridging) atoms on the backbone. This is consistent with the higher electronegativity of the O2 atoms **(Supplementary Fig. S5**). Coupled to the chain compaction observed at 368 K, we observe an increase in the number of Mg^2+^ ions that are bound to O2 atoms of the phosphate groups **(Fig. 2b-c**). Also, the desolvation barrier of Mg^2+^ from the first solvation shell to the second solvation shell around O2 atom is ~2.7 kBT lower at 368 K than the barrier at 298 K **(Supplementary Fig. S5c**). Accordingly, the binding of O2 atoms to a Mg^2+^ ion enables the release of water molecules from the first solvation shell of the Mg^2+^ ion **(Fig. 2d**; **Supplementary Fig. S4**). Loss of water molecules, which is entropically driven at higher temperatures, is compensated by the increased coordination of Mg^2+^ around the polyphosphate. The statistics regarding the radial distribution function of water molecules around Mg^2+^ ions support this idea **(Supplementary Fig. S6**). Mg^2+^ ions can bind to RNA using either outer- (water-mediated contacts) or inner-sphere (direct contacts with the phosphate backbone and nucleobases) interactions. The simulations suggest that raising the temperature results in a shift from outer- to inner-sphere Mg^2+^ contacts. The partially desolvated Mg^2+^ ions can bridge different partially desolvated phosphate groups **(Fig. 2e-f**). This observation is similar to reports of long divalent rod-like ions that act as bridges between like-charge macroions ^64^. We further found that more Mg^2+^ ions are involved in the bridging phosphate groups at higher temperatures **(Fig. 2c**).

**Figure 2:**
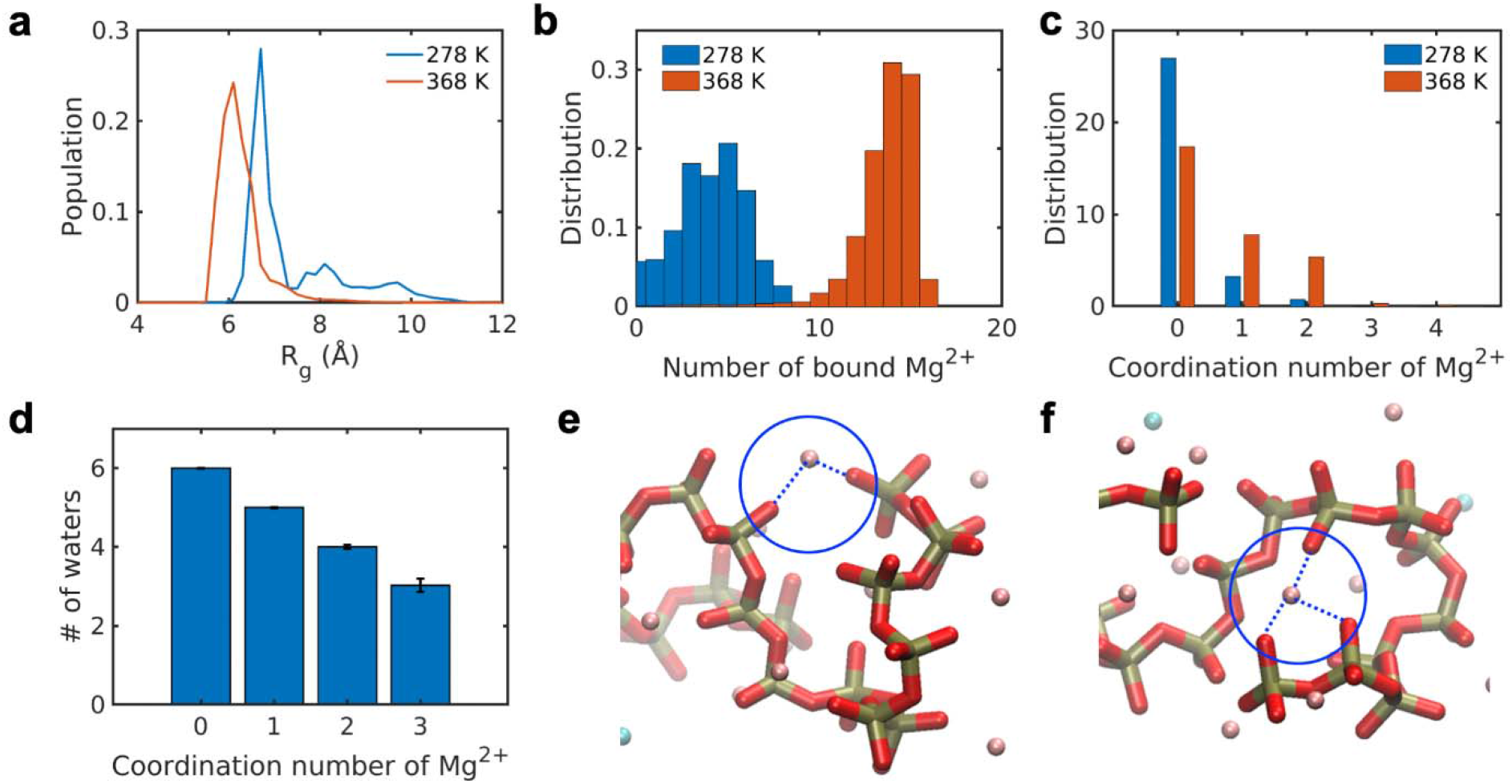
Desolvation and Mg^2+^ ion-induced bridging leads to the compaction of the individual polyphosphate chains. **(a)** Distributions of radii of gyration (R_g_) of polyphosphate at 278 K and 368 K. **(b)** Distribution of number of Mg^2+^ ions bound to O2 (non-bridging) atoms of the polyphosphate at 278 K vs. 368 K. A Mg^2+^ ion is bound to the O2 atom if the distance is smaller than 3 Å, which corresponds to the radius of the first solvation shell of the Mg^2+^ ions around the O2 atom (see **Supplementary Fig. S5a-b**). **(c)** The distribution of coordination numbers of Mg^2+^ at 278 K and 368 K. The coordination number is defined as the number of O2 atoms bound to a Mg^2+^ ion. We observe a shift toward higher coordination numbers, implying a bias toward Mg^2+^ ions acting as bridges between polyphosphate atoms. **(d)** Ensemble-averaged values for the numbers of waters in the first solvation shell around Mg^2+^ ions with different coordination numbers at 368 K. The error bar shows the standard deviation of the distribution of the number of water molecules in the first solvation shell around Mg^2+^ ions. These results show a negative correlation between Mg^2+^ hydration statistics and the coordination numbers around polyphosphate atoms. Snapshots show how a Mg^2+^ ion can be bound to two O2 atoms **(e)** and three O2 atoms **(f)**. In the snapshots, the locations of Mg^2+^ ions are delineated by blue circles.

Chain collapse at higher temperatures, a proxy for phase separation in the single-chain limit ^62, 63^, is entropically driven, originating from the partial desolvation of phosphate groups and Mg^2+^ ions. In the collapsed state, Mg^2+^ ions physically crosslink phosphate groups. Our simulation results show that partial desolvation of phosphate groups and Mg^2+^ ions, coupled to the physical crosslinking role of Mg^2+^ ions, drive the compaction and intra-chain networking of single polyphosphate chains at a higher temperature. These observations are single chain proxies for phase separation coupled to percolation being realizable at higher concentrations. Overall, our simulations suggest that desolvation drives LCST-type phase separation, and the Mg^2+^ mediated physical crosslinks among different chains are likely to engender percolated networks within dense phases.

### LCST coupled to percolation can lead to arrested condensates of RNAs with repeat expansions

Our observation of LCST-type phase transitions for poly(P) and RNA homopolymers suggests that RNAs might have an intrinsic preference for phase separation upon heating in the presence of di- and multivalent salts. This behavior is partly due to the temperature-dependent decrease in free energies of charged groups and polymers ^63, 65, 66^, which for poly(P) leads to partial desolvation of the negatively charged phosphate backbone and the partial desolvation of the Mg^2+^ ions in solution. Based on our findings with homopolymers **(Fig. 1**) and poly(P), we reasoned that RNAs with different nucleobase compositions and thus chemical structure will show variability in LCST phase behavior. To test this idea of sequence-dependent tunability, we first chose to study the effect of temperature on the phase behavior of RNAs with CAG repeats. These RNAs were previously shown to undergo sol-gel transitions *in vitro* ^25, 31, 67^. To examine the phase behaviors of RNAs with CAG repeats, previous studies used an annealing protocol in which the sample was first heated to 95°C and then cooled slowly to 37°C **(Fig. 3a**) ^25, 32, 47^. When imaging was performed immediately after a single heating-cooling ramp, spherical RNA condensates were observed **(Fig. 3a**). We reasoned that the sample containing CAG repeat RNAs formed condensates due to an intrinsic LCST-type phase transition. To test this hypothesis, we used temperature-controlled microscopy (temperature range was 5°C < T < 80°C) to monitor the thermoresponsive transitions of CAGx31 RNA (i.e., 31 CAG repeats).

**Figure 3.**
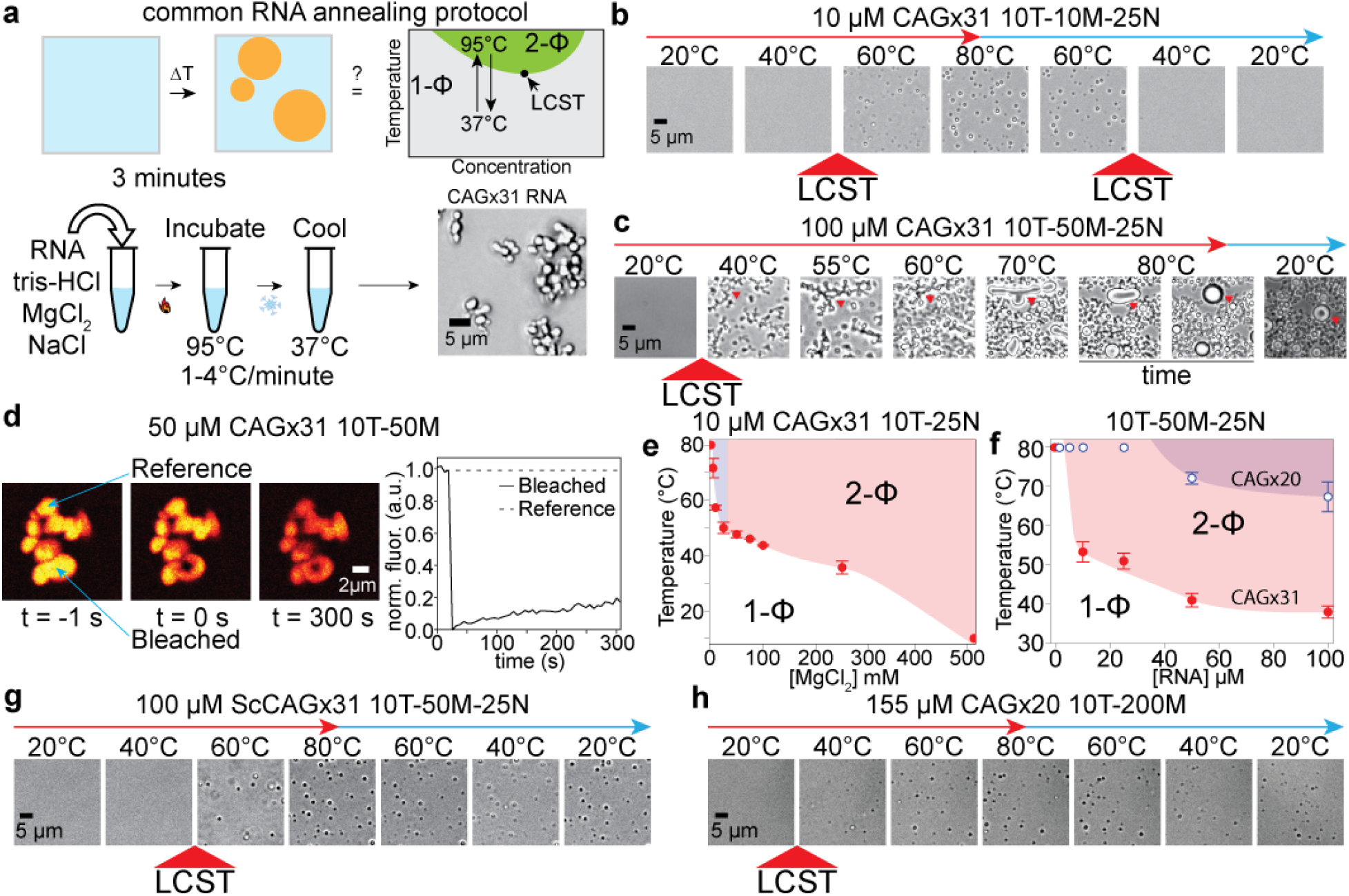
Heat-induced phase separation coupled to percolation of CAG repeat RNAs. **(a)** A schematic of the RNA annealing protocol showing formation of CAGx31 condensates using a brightfield microscope. **(b)** Reversible LCST-type transition of CAGx31 RNA. Brightfield images of 10 μM CAGx31 RNA in 10 mM Tris-HCl (pH 7.5 at 25°C), 10 mM Mg^2+^, 25 mM NaCl (10T-10M-25N buffer); LCPT, 66.8 ± 3.9°C. **(c)** Direct observation of phase separation coupled to percolation of CAGx31 RNA samples. 100 μM of CAGx31 RNA in 10 mM Tris-HCl (pH 7.5 at 25°C), 50 mM Mg^2+^, 25 mM NaCl underwent phase separation at 38.0 ± 3.5°C with aspherical condensates forming immediately at 38.0°C. After settling on the surface, the droplets were subsequently heated gradually to 80°C using a ramp of 1°C/min. During heating, the network of droplets underwent a gradual shape relaxation to spherical condensates. Cooling these spherical condensates did not dissipate them. **(d)** FRAP of CAGx31 condensates. CAGx31 (50 μM unlabeled mixed with 200 nM of Alexa488-labeled RNA) droplets were examined for recovery following photobleaching. **(e)** State diagrams of CAGx31. The cloud point temperatures are plotted as either a function of [RNA] (*left*) or [Mg^2+^] (*right*). Conditions which induced irreversible and reversible phase separation are indicated with red and blue shading, respectively. **(f)** Length dependence of CAG repeat RNA phase separation. State diagrams showing the LCPT as a function of RNA concentration for CAGx31 and CAGx20 (red and blue, respectively). CAGx20 showed an increase in LCPT at comparable concentration as CAGx31. **(g)** Irreversible phase separation of a scrambled variant of CAGx31 that has the same composition as CAGx31 but a different sequence (ScCAGx31; Supplementary Table S1). 50 μM of ScCAGx31 was tested in 10 mM Tris-HCl (pH 7.5 at 25°C), 50 mM Mg^2+^, 25 mM NaCl (10T-50M-25N buffer); LCPT, 57.9 ± 4.2°C. **(h)** Irreversible phase separation of CAGx20. 155 μM CAGx20 RNA was tested in 10 mM Tris-HCl (pH 7.5 at 25°C), 200 mM Mg^2+^ (10T-200M-25N buffer); LCPT, 59.9 ± 2.3°C. Droplets did not dissipate upon cooling. Buffer notation used: the number in front of “T” indicates [Tris-HCl], the number in front of “M” indicates [Mg^2+^], and the number in front of “N” indicates [Na^+^].

Using 10 μM CAGx31 RNA in 25 mM Tris-HCl (pH 7.5 at 25°C), 10 mM MgCl_2_ and 10 mM NaCl (similar to conditions used in an earlier study^25^), we observed a reversible LCST-type transition at 66.8 ± 3.9°C **(Fig. 3b**; **Movie 7**). Increasing the concentration of RNA and Mg^2+^ by five- and 10-fold, respectively, resulted in a significant decrease in the LCST to 38.0 ± 3.5°C **(Fig. 3c**). Under these conditions, we observed that RNA condensation was no longer reversible, and instead, the RNA condensates underwent rapid dynamical arrest **(Supplementary Fig. S7**; **Movie 8**). The arrested RNA condensates persisted even when the sample was cooled well below the lower cloud point temperature (LCPT; **Fig. 3c**; **Supplementary Fig. S4a**; **Movies 9,10**). Upon heating to 30°C above the LCPT, the irregular CAGx31 condensates progressively transformed into spherical liquid-like droplets that showed rapid fusion at high temperatures **(Fig. 3c**; **Supplementary Fig. S8**; **Movies 8,9**). These spherical condensates persisted when subsequently cooled below their LCPT **(Fig 3c**), pointing to hysteresis and dynamical arrest behaviors. Consistent with the idea of dynamical arrest, which results from a hierarchy of intermolecular interactions leading to the formation of a condensate-spanning network, there was minimal recovery of fluorescence after photobleaching at ambient temperature in condensates containing fluorescently labeled CAGx31 **(Fig 3d**). Finally, to rule out any spurious effect of buffer pH change during heating and cooling, we used two additional buffers (HEPES and MOPS) that are known for their lesser sensitivity of pH to temperature than Tris-HCl buffer ^60^. We observed no significant change in the LCPT while titrating Mg^2+^ at a fixed concentration of RNA **(Supplementary Fig. S7b**).

To map the observed thermoresponsive phase separation with higher precision, we generated state diagrams of CAGx31 RNA as a function of varying [RNA] and [Mg^2+^]. Below 10 μM RNA and at 10 mM Mg^2+^, there was no phase separation **(Fig. 3e-f**). When we varied [Mg^2+^] using 10 μM CAGx31 RNA, we found reversibility below 50 mM Mg^2+^ and no phase separation below 1 mM Mg^2+^ **(Fig. 3e**; **Supplementary Fig. S7a**). These data show two distinct temperature- and Mg^2+^-dependent transitions for CAGx31 RNA. The first is an LCST-type phase separation which occurs upon heating and the second is a percolation transition within the dense phase that can cause the dynamical arrest of condensates. Based on our observation of condensate shape relaxation upon heating above LCPT **(Fig. 3c**; **Supplementary Fig. S8**; **Movies 8, 9**), we propose that the percolation transition arises through intermolecular interactions among RNA chains. Thus, while phase separation is favored at temperatures above T_phase_, percolation is disfavored at temperatures higher than T_percolation_ and leads to the melting of networked RNA chains. The dynamical arrest is likely to be determined by whether T_phase_ is higher than or lower than T_percolation_. Irreversibility of condensates or dynamical arrest within condensates is a sign that T_percolation_ is higher than T_phase_.

### Sequence and length dependence of the LCST transition of a GC-rich RNA

To determine whether the LCST-type phase transition was a specific feature of the CAGx31 repeat RNA, we generated a randomized sequence preserving the length and the number of C, A, and G nucleotides with a design constraint that it should not have more than four consecutive G nucleotides to prevent difficulties in RNA synthesis **(Supplementary Table S1**). The scrambled CAGx31 RNA (ScCAGx31) also showed an LCST-type transition. However, the ScCAGx31 RNA showed an elevated T_phase_ **(Fig. 3g**; **Supplementary Fig. S9**) as compared to the CAGx31 repeat RNA under identical experimental conditions (57.9 ± 4.2°C versus 41.1 ± 1.7°C, respectively, for 50 μM RNA in 50 mM Mg^2+^). This observation points to a weakening of the driving forces for LCST phase behavior when the sequence is scrambled. Importantly, the ScCAGx31 condensates became dynamically arrested upon formation **(Movie 10**), akin to what we observed for the CAGx31 RNA. These results suggest that the disruption of precise base-pairing patterns weakens but does not abrogate LCST phase behavior. Next, we performed experiments with CAGx20 RNA (containing only 20 repeats) and a corresponding scrambled RNA (ScCAGx20). Decreasing the RNA length to twenty repeats increased the T_phase_ of the RNA; if we compare 155 μM CAGx20 RNA to 100 μM CAGx31 RNA (to ensure equal numbers of repeats per unit volume), we observe that the T_phase_ increases from 38.0 ± 3.5°C to 59.9 ± 2.3°C **(Fig. 3h**; **Movie 11**). Interestingly, ScCAGx20 showed a substantially lower LCPT compared to CAGx20 **(Fig. 3h**; **Supplementary Fig. S9b**). This scrambled 60-mer RNA had an LCPT that was comparable to CAGx31.These observations point to an intriguing interplay of backbone-mediated and nucleobase-specific entropic contributions arising from desolvation, contributions from configurational entropy as evidenced by the lowering of LCPT with increased chain length, and the contributions of base pairing / stacking interactions that are different in scrambled versus repeat sequences of different lengths.

### Uracil suppresses LCST phase transitions

To further assess the contribution of RNA sequence on the phase separation properties of repeat RNAs, we sought to compare CUG and CUU repeats with CAG repeat RNAs. The choice of these two repeat RNAs was inspired by our experiments with homopolymeric RNAs, which revealed that poly(rU) RNA displays a UCST-type transition in the presence of Mg^2+^ without any detectable LCST transition between 5 and 80°C **(Fig. 1**). Based on these results, we hypothesized that uracil may act to suppress LCST-type phase transitions in repeat RNAs **(Fig. 4a**). To test this postulate, we generated two variants of the CAGx31 sequence, one with an A-to-U substitution (CUGx31) and the other with an A-to-U and a G-to-U substitution (CUUx31). Using temperature-controlled microscopy, we found that CUGx31 RNA exhibited an LCST-type phase transition across a range of RNA and Mg^2+^ concentrations as was observed with the CAGx31 RNA. However, the LCPT of CUGx31 RNA was substantially higher compared to CAGx31 RNA under similar conditions (52.8 ± 1.2°C vs. 41.1 ± 1.7°C, 50 μM RNA in 50 mM Mg^2+^; **Fig. 4b**). These data suggest that the A-to-U substitution weakens phase separation. Interestingly, while CAGx31 RNA formed irreversible dynamically arrested condensates, CUGx31 RNA condensates were reversible under identical conditions [10 mM Tris-HCl (pH 7.5 at 25°C), 50 mM Mg^2+^, 25 mM NaCl; **Fig. 4c**; **Movie 12**]. Finally, unlike CAGx31 and CUGx31 RNAs, we found that CUUx31 RNA did not undergo phase separation under all conditions tested **(Fig 4d**; **Movie 13**). These results collectively support the idea that RNA phase separation coupled to percolation can be tuned by the nucleobase composition, with uracil weakening LCST-type transitions as well as abolishing the hallmarks of dynamical arrest such as hysteretic phase behavior and percolation.

**Figure 4.**
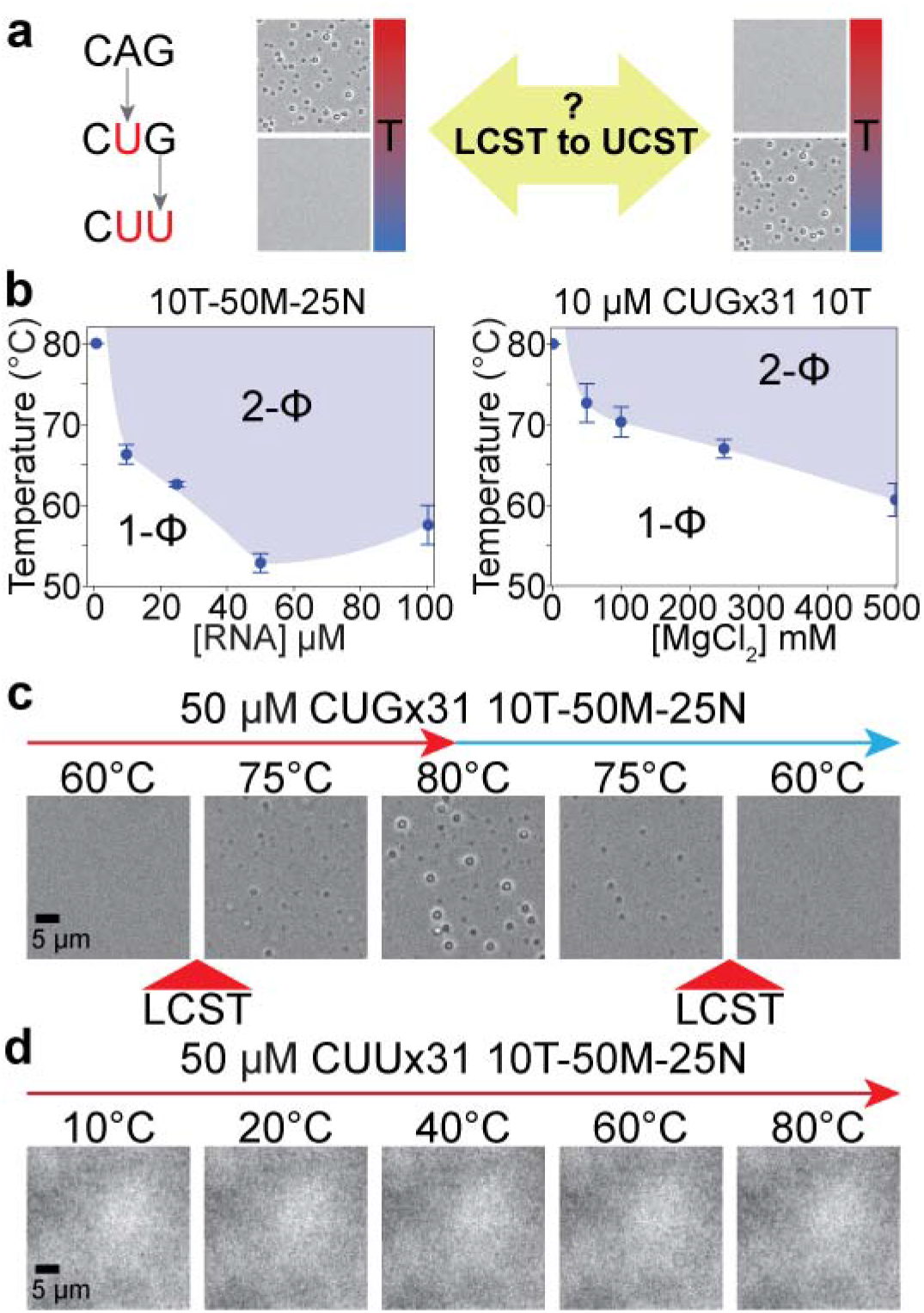
Uracil suppresses LCST-type phase separation of repeat RNAs. **(a)** A schematic of our repeat RNA sequence design. **(b)** State diagrams of CUGx31. Shown here is a plot revealing the dependence of the LCPT of CUGx31 on RNA concentration in 10 mM Tris-HCl (pH 7.5 at 25°C), 50 mM Mg^2+^, 25 mM NaCl (10T-50M-25N buffer) (*left*). The dependence of the LCPT of 10 μM CUGx31 on the concentration of Mg^2+^ is also shown (*right*). **(c)** Reversible condensation of CUGx31 RNA upon heating. Brightfield images of 50 μM CUGx31 RNA in 10 mM Tris-HCl (pH 7.5 at 25°C), 50 mM Mg^2+^ and 25 mM NaCl (10T-50M-25N buffer) that was subjected to heating (red arrow) and cooling (cyan arrow) as indicated; LCPT, 72.6 ± 5.4°C. **(d)** Brightfield images of 50 μM CUUx31 RNA during heating (same buffer conditions as panel **c**); no phase separation was observed in this case. Buffer notation used: the number in front of “T” indicates [Tris-HCl], the number in front of “M” indicates [Mg^2+^], and the number in front of “N” indicates [Na^+^].

### Catalytic RNAs display rich thermoresponsive phase separation properties

To test the generality of RNA LCST-type phase separation, we examined longer RNAs with higher sequence complexity. We focused on RNase P, a Mg^2+^-dependent endonuclease that catalyzes tRNA 5’ maturation in all three domains of life. In the RNP form of RNase P, a single catalytic RNase P RNA (RPR; **Fig. 5a**) works together with protein co-factors to perform tRNA processing *in vivo* ^68^.^69^ However, since RPRs alone exhibit multiple turnover during pre-tRNA cleavage *in vitro* ^70^, RPRs offer a robust model system to assess the effect of RNA PSCP on RNA function. Here, we investigated archaeal RPRs from two thermophiles [*Pyrococcus furiosus* (*Pfu*) and *Methanocaldococcus jannaschii* (*Mja*)] and a mesophile [*Methanococcus maripaludus* (*Mma*)]. While RNase P from these three archaea have been extensively characterized ^68, 71, 72, 73, 74, 75^, the *Pfu* RPR is of special interest given that its activity in the absence of protein cofactors has been studied under different conditions ^74, 76^.

**Figure 5.**
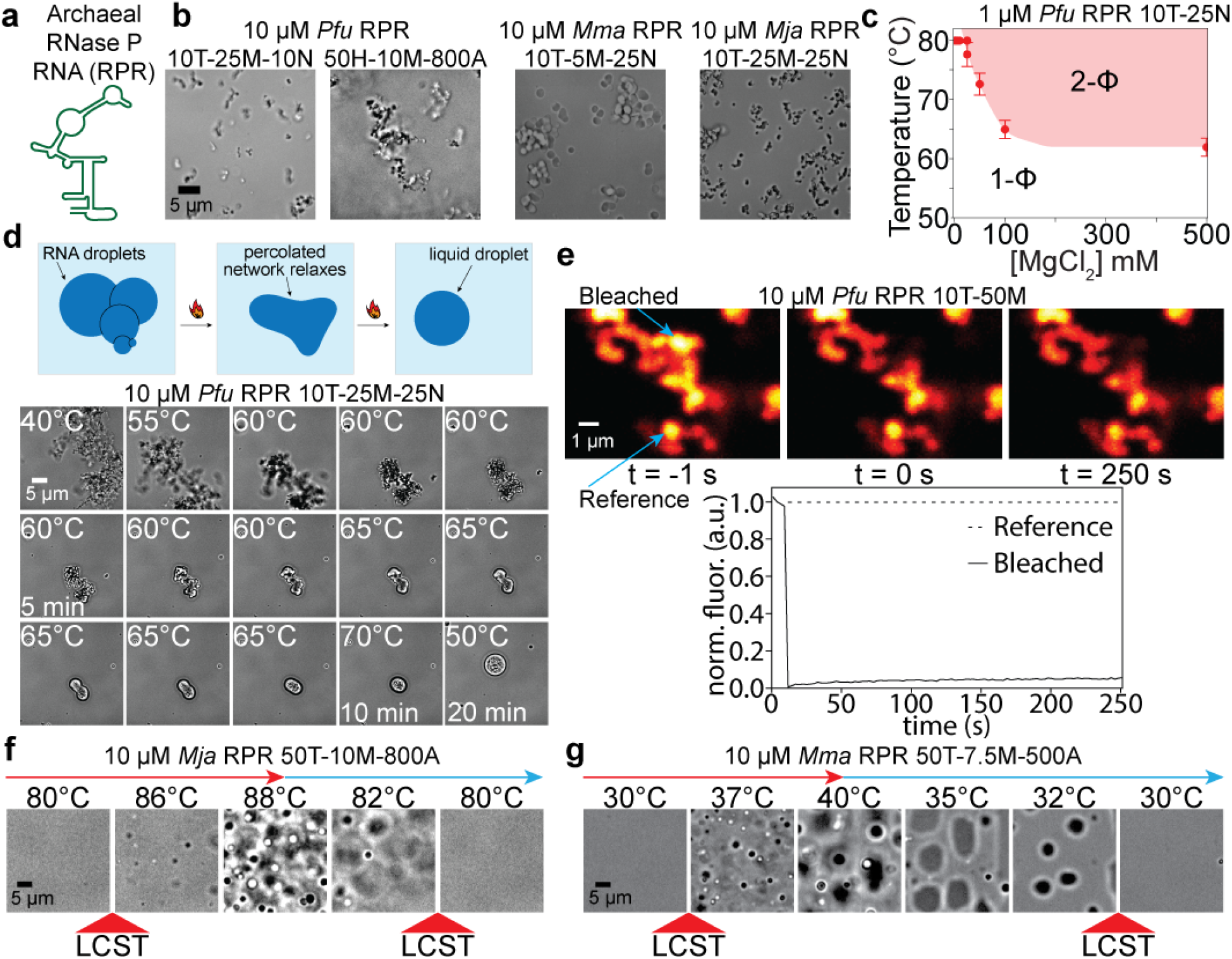
Phase behavior of RNase P RNAs. **(a)** Cartoon representation of the RNase P RNA (RPR). **(b)** *Pfu, Mma*, and *Mja* RPRs form condensates after a single heating-cooling ramp. The buffer conditions were 10 mM Tris-HCl (pH 7.5 at 25°C); 10 mM, 5 mM or 25 mM Mg^2+^ for *Pfu*, *Mma*, and *Mja*, respectively; and 25 mM NaCl. *Pfu* refolding buffer contains 50 mM HEPES-KOH (pH 8.5 at 25°C), 10 mM Mg^2+^, and 800 mM AmAc (50H-10M-800A). **(c)** A state diagram of *Pfu* RPR. Shown here are LCPTs for various concentrations of Mg^2+^ ions. **(d)** Heat-induced shape relaxation of arrested RNA condensates (*left-to-right). Pfu* RPR (1 μM) was subjected to an annealing cycle and the resultant droplet clusters were imaged in the same buffer as that used for annealing in **b** [10 mM Tris-HCl (pH 7.5 at 25°C), 25 mM Mg^2+^, 25 mM NaCl]. **(e)** FRAP of *Pfu* RPR condensate clusters. Condensates were prepared by annealing 10 μM fluorescein-labeled *Pfu* in a buffer containing 10 mM Tris-HCl (pH 7.5 at 25°C), 50 mM Mg^2+^. **(f)** *Mja* RPR demonstrated reversible phase separation in refolding buffer containing 50 mM HEPES-KOH (pH 8 at 25°C), 10 mM Mg^2+^, and 800 mM AmAc. **(g)** *Mma* RPR demonstrated reversible phase separation in refolding buffer containing 50 mM Tris-HCl (pH 7.5 at 25°C), 7.5 mM Mg^2+^, and 500 mM AmAc. Buffer notation used: the number in front of “T” indicates [Tris-HCl], the number in front of “H” indicates [HEPES-KOH], the number in front of “M” indicates [Mg^2+^], the number in front of “N” indicates [Na^+^], and the number in front of “A” indicates [AmAc] in a given buffer.

We initially prepared RPRs following the annealing protocol used in an earlier study that examined repeat RNAs^25^ **(Fig. 5b**). We tested a range of Mg^2+^ and Na^+^ concentrations as well as buffer conditions specifically designed to promote optimal RPR folding ^68, 71, 72, 73, 74, 75^. With *Pfu, Mja*, and *Mma* RPRs, we observed Mg^2+^-dependent formation of condensates **(Fig. 5b**). When [Na^+^] was varied at a fixed [Mg^2+^], we observed distinct morphological changes in the RPR condensates **(Supplementary Fig. S10**). While condensates were small and spherical at low and high NaCl concentrations, we found that for the *Mja* RPR concentrations of 100 to 250 mM NaCl resulted in condensates with arrested coalescence ^77^ **(Supplementary Fig. S10c**). However, irrespective of their appearance, all RPRs were observed to display a clear LCST-type phase separation followed by a percolation transition similar to the CAG repeat RNAs **(Fig. 5b-c**; **Supplementary Fig. S10**; **Movies 14-16**). Moreover, the percolated RPR condensates underwent shape relaxation when heated above their LCPT **(Fig. 5d**; **Movie 17**), suggesting that intermolecular interactions among RNA chains are the drivers of dynamic arrest. Consistent with this interpretation, we observed that FTSC-labeled RPR condensates displayed little fluorescence recovery upon photobleaching at room temperature **(Fig. 5e**; **Supplementary Fig. S11**; **Movies 18-20**).

Most laboratories have customized folding protocols for their RNAs of interest ^50, 76, 78, 79, 80^; in this regard, RPRs are no exception ^71, 72, 73, 74^. RNA folding is hierarchical with nascent secondary structures transitioning through discrete steps to generate a tertiary fold. Folding protocols are empirically optimized for a combination of mono/divalent cations that facilitate the attainment of a functional 3-D structure. Since percolated RNA condensates will deplete soluble RNA molecules, we hypothesized that previously optimized buffer and annealing conditions ^71, 72, 73, 74^ (termed as refolding buffer) for RPR folding may suppress phase separation and percolation, the latter being the determinant of condensate arrest. To test this idea, we performed heating and cooling of *Pfu*, *Mja*, and *Mma* RPRs in their respective refolding buffers, all of which contain ammonium acetate (see *Materials and Methods*) and monitored mesoscale changes continuously. Remarkably, while *Mma* and *Mja* RPRs formed reversible condensates under these conditions **(Fig. 5f-g**; **Movies 21, 22**), *Pfu* RPR did not show any visible phase separation **(Movie 23**). We finally tested the effect of ammonium acetate (AmAc) alone, which is commonly used in RNA folding regimens and previously observed to suppress RNA sol-gel transition ^25^. Indeed, ammonium acetate abrogated the LCST-type transition of RPR RNAs **(Supplementary Fig. S12**; **Movies 14, 24**).

### Phase separation of RNA inhibits RPR enzymatic activity in vitro

Peptide-mediated phase separation has previously been shown to modulate ribozyme activity ^81, 82^. However, it is unclear how protein-independent ribozyme phase separation might alter its enzymatic function. To determine the effect of PSCP on *Pfu* RPR activity, we measured its pre-tRNA cleavage activity after refolding it under conditions that either suppress (+AmAc) or promote (-AmAc) PSCP **(Supplementary Fig. S12**; **Movies 23, 24**). Our approach was as follows: at 10 mM and 50 mM Mg^2+^, *Pfu* RPR exhibits a T_phase_ of 81°C and 69°C, respectively. We therefore chose to assay the RPR at temperatures 5°C above or below the T_phase_. Our strategy was intended to assess if activity-altering effects were more likely above rather than below the T_phase_. Subjecting the *Pfu* RPR to a 50°C annealing protocol that includes 800 mM AmAc (our standard procedure^74^) yields activity that is comparable to the RPR annealed at 76°C or 86°C (10 mM Mg^2+^; **Fig. 6b *left***) or at 64°C or 74°C (50 mM Mg^2+^; **Fig. 6b *right***) in presence of 800 mM AmAc. This finding suggests that incubating the RPR at high temperatures under conditions that suppress condensate formation does not significantly dampen function. In sharp contrast, there is at least a 4- to 6-fold decrease in *Pfu* RPR activity on account of phase separation into arrested condensates, an outcome of annealing at temperatures above the T_phase_ and in the absence of AmAc **(Fig. 6b**).

**Figure 6:**
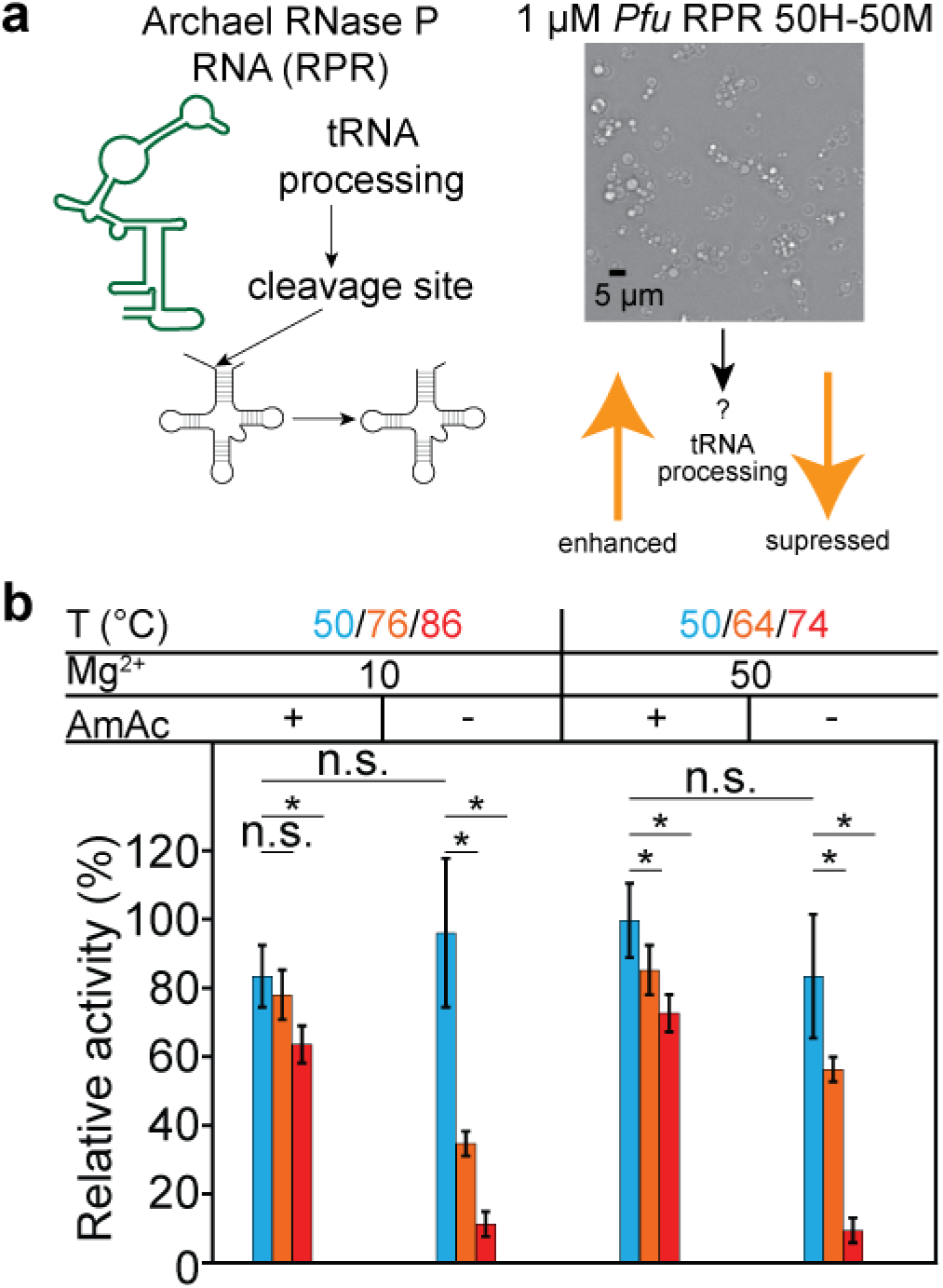
Phase separation inhibits the activity of *Pfu* RNase P RNA. **(a)** Schematic of pre-tRNA cleavage catalyzed by RPRs. **(b)** The pre-tRNA cleavage activity of *Pfu* RPR was measured under a range of conditions that suppress or promote phase separation. *Pfu* RPR (2.5 μM) was annealed in buffer containing 50 mM HEPES-KOH, pH 7.5 (55°C) and the specified concentrations of MgCl_2_ (10 or 50 mM) and AmAc (0 or 800 mM) and at the indicated temperatures for 5 min. After annealing, RPRs were tested for their ability to cleave *Eco* pre-tRNA^Tyr^ (30 μM). The average turnover numbers, which were normalized according to the highest measured cleavage rate (50 mM MgCl_2_, 800 mM AmAc, 50°C), and their associated standard deviations were determined from at least three technical replicates. Buffer notation used: the number in front of “H” indicates [HEPES-KOH] and the number in front of “M” indicates [Mg^2+^] in each buffer.

To verify that the observed loss of activity was indeed due to RNA condensate formation and not due to Mg^2+^-dependent RPR degradation at high temperatures, we analyzed annealed RPRs by agarose gel electrophoresis **(Supplementary Fig. S13**). Consistent with our observation that *Pfu* RPR retains a high level of activity when annealed at 50°C, a single species with expected mobility was observed after annealing at 50°C under all combinations of Mg^2+^ and AmAc concentrations tested. In contrast, under conditions that induce phase separation (0 mM AmAc, T_phase_+5°C), we observed that the RPR was trapped in the well suggesting the presence of high molecular weight species that were unable to enter the gel. Taken together with the optical imaging and FRAP data **(Fig. 5b, e**), we conclude that the arrested condensates contain the RPR but not in a physical form that is resolved by gel electrophoresis.

## Discussion and Conclusions

Protein-independent phase transitions of RNAs are garnering interest due to intracellular condensation of trinucleotide-repeat expansion RNAs associated with neurodegenerative disorders ^25^, implications for RNA catalysts in the prebiotic scenario ^83^, interests in creating RNA-based artificial cells ^84, 85^, and the aspiration to design programmable nucleic acid-based soft-matter ^58, 85, 86, 87^. Here, we performed experiments to shed light on the origin of RNA phase transition and the role of RNA-RNA interactions in this process. Our results suggest that RNA chains have an intrinsic ability to undergo an LCST-type phase transition that is driven by entropy gain due to chain desolvation ^88, 89^, enhanced divalent metal ion binding at elevated temperatures ^90^, and ion-induced bridging interactions between RNA chains ^91, 92, 93^. In contrast, the RNA-RNA interactions via base pairing and stacking ^25, 47^ drive percolation transitions within condensates formed by LCST-driven phase separation. Our experiments with inorganic poly(P) suggest that the LCST-type phase transition is a generic property encoded by the negatively charged phosphate backbone of RNA, with positive or negative effects imparted by the specific RNA sequence composition. While poly(rC) and poly(rA) showed a clear LCST-type transition in the presence of Mg^2+^ ions, poly(rU) exhibited only a UCST-type transition under similar conditions. Nevertheless, poly(rU) chains were previously shown to have access to both UCST and LCST-type transitions in the presence of multivalent ions ^22^. Further, recent studies have shown that in monovalent salts, and in the presence of polyethylene glycol, RNA homopolymers exhibit UCST phase behavior ^94^. Therefore, segregative transitions, i.e., phase separation, can be either LCST- or UCST-type depending on the solution conditions and types of solution ions, which directly influence the temperature dependent free energies of solvation. As a result, complex closed-loop or hour-glass type phase diagrams are the most likely scenario for RNA molecules ^22^. These are likely to be unmasked under specific sets of conditions or alternatively, as shown in this work, the specific choice of solution conditions amplifies one type of segregative transition. As for the associative, percolation transitions, they are likely to always be enthalpically driven.

Collectively, our results lead to the following model for RNA LCST behavior. For a given [Mg^2+^] and [RNA], there is a hierarchy of LCSTs: poly(rU) (not measurable under our experimental conditions) > poly(rC) > poly(P) > poly(rA) > poly(rG) **(Fig. 7a**). The implication is that the LCST of poly-purines will be lower than poly(P) whereas the LCST of poly-pyrimidines will be higher than that of poly(P). Purines extract a higher entropic penalty for being solvated and hence the LCST is lowered vis-à-vis poly(P). Likewise, the pyrimidines extract a lower entropic penalty as compared to poly(P), thus raising the LCST. As for the comparisons between rG and rA, the former, in the context of base pairing and hydrogen bonding in general has at least two donors and one acceptor whereas rA has one donor and one acceptor. The extra functional group extracts a higher entropic penalty for organizing solvent molecules, implying lowering of LCST. Likewise, rC, in the context of hydrogen bonding has at least one donor and two acceptors whereas rU has one of each.

**Figure 7.**
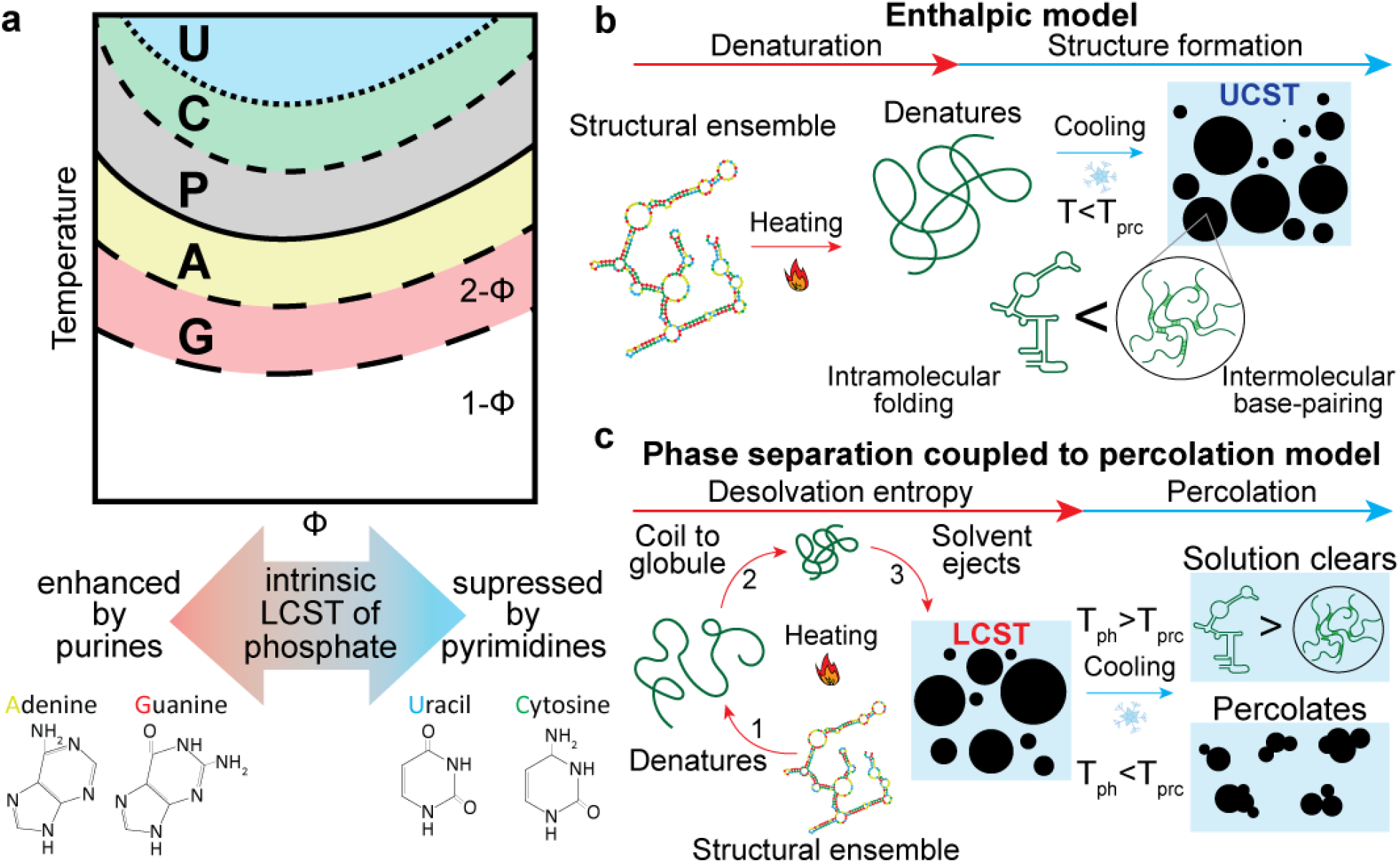
An emerging model for RNA phase separation coupled percolation (PSCP) behavior. **(a)** The observed hierarchy of LCST-type phase transition propensity of RNA bases and the phosphate backbone. **(b)** An enthalpic model of RNA phase separation. Here, RNA in an ensemble of minimum free energy structures (*left*) is heated, thereby denaturing the RNA. Subsequent cooling should enable the RNA to undergo a UCST transition. This model implies that enthalpic interactions such as hydrogen bonding and base-stacking drive RNA phase separation. **(c)** PSCP model of RNA condensation. In this model, desolvation entropy drives the self-association of RNA to form phase-separated condensates upon heating. During subsequent cooling, the rank order of the phase separation temperature (LCPT or T_ph_) and the percolation temperature (T_prc_) determines refolding *vs*. condensate arrest. Our experimental and computational results clearly show that RNA condensation proceeds via this pathway.

Upon phase separation, RNA-RNA interactions in the dense phase can lead to the formation of a percolated network. Since base-pairing and stacking interactions are temperature-dependent and disrupted at higher temperatures ^95, 96^, the RNA condensates transition to a predominantly viscous state at temperatures higher than the characteristic “percolation temperature” of the network (T_prc_). This observation imposes an orthogonal temperature response of percolation and RNA phase transitions **(Fig. 7c**). Based on our results, we propose a conceptual model that sheds light on the formation of previously reported RNA condensates through a single heating-cooling ramp *in vitro* ^23, 25, 31, 32, 36^. An important question is why do these condensates remain spherical after their preparation but behave as static aggregates without the characteristic dynamicity of phase-separated liquids? To answer this question, we consider the relative rank order of lower cloud point temperature (LCPT) and percolation temperature (T_prc_) in our model. If the LCPT is lower than the percolation temperature, the RNA chains are arrested in the phase separated state. If T_prc_ is lower than the LCPT, reversible RNA phase separation is realized. The persistence of spherical RNA condensates well below their LCPT is a hallmark of condensates formed by LCST-type phase separation coupled to percolation (PSCP; with T_prc_ > LCPT). Importantly, the phase separation and percolation transition temperatures and their relative rank order are dependent on the buffer composition, RNA chain length, sequence composition, and nucleobase patterning. This idea is exemplified by our contrasting results with CAG versus CUG versus CUU repeat RNAs **(Figs. 3, 4**). CAGx31 RNA undergoes an LCST-type transition when heated and the condensates subsequently form a percolated system-spanning network when cooled, giving rise to arrested RNA condensates at Mg^2+^ concentrations > 25 mM. In contrast, CUGx31 undergoes reversible LCST-type phase separation and CUUx31 RNA remained soluble under similar conditions.

We consider a few important implications of our findings. Firstly, the ubiquitous ability of RNAs to undergo LCST-type phase separation is highlighted by our experiments with RPR RNAs featuring longer chain lengths and higher sequence complexity as compared to the trinucleotide repeat sequences. Leveraging the *Pfu* RPR ribozyme ^76^, we assessed the effect of PSCP on the activity of this catalytic RNA. Our observations suggest that the dynamical arrest of RPR chains within phase separated condensates inhibits RPR catalysis. These findings are relevant to the RNA world hypothesis ^97^ which postulates that the early crucible of life leveraged RNA both as the genetic material and as the workhorses that catalyzed prebiotic metabolic reactions. Condensates could engender payoffs for RNA catalysis through mass action effects or inducing a switch from a less to a more active enzyme conformation, thus reversing the fate of ostensibly unfavorable reactions. Nevertheless, such benefits are moot if the catalyst is devoid of functional structure and is entangled in a network, as evident from the decrease in activity upon dynamical arrest of *Pfu* RPR chains within phase-separated condensates **(Fig. 6**). Given the tunability of the percolated network by the target RNA’s make-up and buffer conditions, however, we recognize that condensates could either accelerate or dampen catalysis. In fact, both increased and decreased rates have been reported for compartmentalized RNA (hammerhead ribozyme) catalysis ^82, 98^. Thus, the evolution of innovations pertaining to the diverse biological roles of RNAs must have entailed balancing the potential kinetic benefits of phase separation versus the potential loss of function associated with ensnarement in an irreversible, percolated network.

Second, we observed that the arrested RNAs can be recovered by condensate reversal through the treatment of EDTA, which sequesters Mg^2+^ ions from RNA chains **(Supplementary Fig. S14**). This observation points to an intriguing possibility that RNA LCST-type phase separation provides a spontaneous physical process for reversible storage of RNA molecules under heat stress. Moreover, the facile reversibility afforded by the potential sequestration of Mg^2+^ by key metabolites (e.g., citrate ^99^) suggests an exciting nexus between cellular metabolic state and RNA phase separation/function.

Third, from a physicochemical perspective, percolation transition can be inhibited by factors/additives that can modulate RNA base-pairing and stacking interactions. For example, while phase-separated CAG repeat RNAs are dynamically arrested *in vitro*, Jain and Vale reported that the CAG granules were liquid-like *in cellulo* ^25^. They also posited that helicases, which are ATP-dependent RNA binding proteins, can remodel RNA-RNA interactions within RNA granules thereby giving rise to the observed fluidity of CAG-repeat condensates in cells. Considering our model formulated based on our results **(Fig. 7c**), we propose a testable idea that helicases may act as suppressors or modulators of condensate percolation without necessarily altering the RNA LCST-type phase separation. The ability of proteins to alter a RNA’s LCST-type phase separation and/or percolation characteristics may provide a new lens to view the roles of proteins in RNP granule formation and regulation.

From a practical standpoint, our findings should foster a new approach to optimization of laboratory procedures that involve heating and cooling of nucleic acids (e.g., PCR, fluorescence *in situ* hybridization, RNA folding, nucleic acid synthesis/purification)^100, 101, 102, 103^. In many of these experiments, an additive (e.g., ammonium acetate ^25, 104^, DMSO ^105^) is empirically determined to increase the process efficiency. Our experimental results indicate that such additives can modulate the LCST-type phase transitions of RNA as well as the accompanying percolation. Specifically, we observed that ammonium acetate suppresses phase separation while DMSO enhances RNA condensation **(Supplementary Fig. S15**). Although such optimization steps are typically performed in an *ad hoc* fashion, our results should encourage a more rational approach to examine the phase behavior of a nucleic acid of interest under different buffer conditions.

In summary, we present phase separation coupled to percolation as a general mechanism for RNA condensation based on the rank order of the LCPT and the percolation temperature. Our results indicate that desolvation entropy plays a major role in the phase separation of RNA chains while enthalpy drives the percolation transition. Our results raise important questions about the role of RNAs in the thermoresponsive phase behavior of ribonucleoprotein (RNP) granules, and how RNP binding alters RNA LCST-type phase separation and/or percolation. Finally, RNA phase separation at higher temperatures may be functionally relevant for RNA catalysis in organisms that thrive under extreme conditions such as psychrophiles and thermophiles.

## Authors’ Contributions

P.R.B. conceived the idea of this study. P.R.B. and G.M.W. designed the study with input from L.B.L, W.J.Z., V.G., X.Z., and R.V.P. G.M.W performed the RNA phase separation experiments and data analysis with assistance from P.P. L.B.L, V.S. and W.J.Z. synthesized the different RNAs and modified them with fluorescent labeling. W.J.Z. performed all the RNase P activity experiments. X.Z. and R.V.P. performed all-atom MD simulations, analyzed the results, and developed the framework to explain the observed phenomenology. All authors contributed in writing and revising the manuscript.

## Acknowledgments

This work was supported by grants from the National Institute of General Medical Sciences of the National Institutes of Health (R35 GM138186 to P.R.B, GM120582 to VG), the Air Force Office of Scientific Research (grant FA9550-20-1-0241 to R.V.P), and the St. Jude Research Collaborative on Biophysics of RNP granules (to P.R.B and R.V.P). WJZ gratefully acknowledges a Pelotonia postdoctoral fellowship from the OSU Comprehensive Cancer Center. The authors acknowledge members of the Banerjee, Gopalan, and Pappu labs for valuable discussions during different stages of the manuscript preparation. X.Z. thanks Alan A. Chen and François-Yves Dupradeau for helpful discussions about the forcefield parameters used in this work, and Stephen Tahan for support in the using of RIS cluster at Washington University in St. Louis.

