## Supplementary Information for "RNAs undergo phase transitions with lower critical solution temperatures"

[§]These authors contributed equally

*Correspondence should be addressed to:

### Materials and Methods

#### In vitro transcription (IVT) and purification of RNA

Unless specified as commercially synthesized, all RNAs were generated by run-off IVT using T7 RNA polymerase. The different archaeal RPRs used in this study were previously cloned in pBT7[1] and each placed under the control of a T7 RNA polymerase promoter. The DNA templates for IVT of *Pfu, Mja, and Mma* RPRs were prepared by PCR using as starting material pBT7-*Pfu* RPR[2], pBT7-*Mja* RPR[3], and pBT7-*Mma* RPR [4]. The *Pfu* RPR template was generated by PCR-based amplification using the forward primer F-ext and the reverse primer R-ext, and subsequent digestion of the resulting PCR amplicon with *Eco*RI. Anticipating some future studies that entailed annealing a fluor-conjugated oligo, both *Mja* and *Mma* RPRs were synthesized as variants containing 23-nt 5'- and 3'-extensions. To accomplish this goal, we used a two-step PCR. The *Mja* and *Mma* RPR templates were generated by first amplifying the sequences using the appropriate forward and reverse primers (*Mja*5ext-F and *Mja*3ext; *Mma*5ext-F and *Mma*3ext-R). The resulting PCR products were then amplified again using the forward primer T7ext-F and the respective reverse primers. Sequences of all oligonucleotide primers used in this study are listed in **Supplementary Table S1**.

To prepare the IVT templates for generating the CAG RNAs, constructs with a double-stranded T7 RNA polymerase promoter followed by a single-stranded coding sequence were obtained by annealing T7pro-sense to the appropriate antisense (AS) oligo (**Supplementary Table S1**) in a 3:8 mass ratio.

A 100 µL IVT reaction contained template (0.4 µg for the RPRs and 11 µg for the CAG RNAs); 1x IVT buffer [40 mM Tris-HCl (pH 7.6 at 22°C); 24 mM $MgCl_2$; 2 mM spermidine; 0.01% (v/v) Triton X-100]; 10 mM DTT; 5 mM each of ATP, CTP, GTP, and UTP (UTP was omitted for the CAG RNAs); 0.002 U thermostable inorganic pyrophosphatase (NEB); and T7 RNA polymerase (purified in-house). After a ~5-h incubation at 37°C, the reactions were treated with DNase I (Roche), extracted with phenol-chloroform, and dialyzed using 3,500-MWCO tubing against 3 L $ddH_2O$ (three changes in 20 h) at 4°C. The RNAs were then precipitated with 0.3 M sodium acetate and 2.5 volumes of ethanol, resuspended in autoclaved $ddH_2O$, and quantitated by measuring the absorbance at 260 nm and using the respective extinction coefficients (OligoAnalyzer Tool, IDT).

*Pfu* RPR, *Mja* RPR, *Mma* RPR, CAGx20, ScCAGx20, CAGx31, and ScCAGx31 (500 µg each) were 3' labeled with fluorescein thiosemicarbazide (FTSC; kind gift of Prof. Edward Behrman, OSU). RNAs were first incubated with a 100-fold molar excess of sodium periodate and 100 mM sodium acetate (pH 5.2 at 22°C) for 90 min at 25°C in the dark. The RNAs were then precipitated with 2.5 volumes of ethanol and resuspended in autoclaved $ddH_2O$. The oxidized RNAs were then incubated with a 30-fold molar excess of FTSC and 100 mM sodium acetate (pH 5.2 at 22°C) for ~16 h at 25°C in the dark. The labeled RNAs were then precipitated with 2.5 volumes of ethanol and re-suspended in autoclaved $ddH_2O$. The fluor-labeled RNAs were loaded onto a 6% (w/v) polyacrylamide gel containing 7 M urea. After UV shadowing, the RNAs of interest were extracted and eluted in 100 mM sodium acetate (pH 5.2 at 22°C) and 5 mM EDTA first for 3 h at 37°C and then overnight at 4°C. The eluted RNAs were precipitated with 2.5 volumes of ethanol, re-suspended in autoclaved $ddH_2O$, and quantitated by measuring the absorbance at 260 nm and using the respective extinction coefficients. The purity of the gel-extracted RNAs was assessed by electrophoresis on an 8% (w/v) polyacrylamide gel containing 7 M urea and imaging the gel using the Cy2 setting on an Amersham Typhoon RGB (Cytiva).

#### RNA sample preparation via annealing method

RNAs that were purchased from Integrated DNA Technologies (see **Supplementary Table S1**) were resuspended in RNase free water to a final concentration of 200 µM and stored at −20°C.

RNA samples that were prepared by IVT (see **Supplementary Table S1**) were stored in RNase free water at −20°C. Appropriate buffer stocks were prepared by mixing 1 M Tris-HCl (pH 7.5 at 25°C) or 1 M HEPES-KOH (pH 7.5 at 25°C); 5 M NaCl; and 1 M $MgCl_2$ stock solutions in RNase free water. RNA was added to the buffer as the last ingredient at a final working concentration of 10 µM or as specified in the text. A 5 µl sample was added to a 0.65-mL microcentrifuge tube and placed on a heating block preheated to 95°C. After 3 min, the heat block was turned off and the RNA samples were allowed to slowly cool to 37°C over ~5 h. This protocol is similar to previously reported methods [5, 6, 7, 8, 9]. A 2 µl sample was then placed on a Tween-20 coated glass slide for brightfield imaging of condensates using a Zeiss Primovert microscope equipped with a 40X air objective and Zeiss Axiocam.

#### RNase P activity assay

*Pfu* RPR (2.5 µM) was first diluted in annealing buffer containing 50 mM HEPES-KOH (pH 7.5 at 55°C), 10 or 50 mM $MgCl_2$, and 0 or 800 mM $NH_4OAc$. Addition of $NH_4OAc$ suppresses phase separation. RPR in annealing buffer was incubated for 5 min at ±5°C of $T_{phase}$, which was determined through microscopy methods to be either 81°C or 69°C in the presence of 10 mM or 50 mM $MgCl_2$, respectively, and then subsequently cooled to room temperature (19–21°C). In addition, to test whether annealing temperature affects RPR cleavage activity even without phase separation, we performed a control experiment in which the *Pfu* RPR in annealing buffer was incubated at 50°C for 5 min before cooling to room temperature.

To initiate the cleavage reaction, RPR in annealing buffer was incubated at 55°C for 10 min before adding *Eco* pTyr substrate (30 µM), a trace amount of which was 5′ γ-$^{32}$P radiolabeled. Upon addition of substrate, the final reaction buffer was adjusted to 60 mM HEPES-KOH (pH 7.4 at 55°C); 2 M $NH_4OAc$, and 500 mM $MgCl_2$, and the final concentration of *Pfu* RPR was 0.625 µM. At pre-defined time points, aliquots of the reaction mix were quenched in stop dye containing 7 M urea, 1 mM EDTA, 10% (v/v) phenol, 0.04% (w/v) bromophenol blue, and 0.04% (w/v) xylene cyanol. Quenched reaction aliquots were separated on a 10% (w/v) polyacrylamide gel containing 7 M urea. Gels were exposed to a storage phosphor screen that was subsequently scanned using an Amersham Typhoon RGB phosphorimager (Cytiva). Bands corresponding to uncleaved and cleaved pTyr were quantitated using ImageQuant (Cytiva) and the data were plotted and fitted to a straight line using KaleidaGraph (Synergy Software) to determine reaction velocities and turnover numbers. Significance was determined using p-values calculated using the pandas and scipy libraries in python. The reported turnover numbers and error bars were determined from at least three technical replicates.

#### Sample preparation for temperature-controlled microscopy

Borosilicate glass slides (1” x 3”) were cut to be approximately 20 mm x 25.4 mm using a carbide blade glass scribe (Thorlabs). Slides and 18 mm square #1.5 coverslips were coated with Tween-20 by incubating them in a coplin jar filled with a 20% (v/v) solution of Tween-20 and MilliQ water for 30 min, and then rinsing five times in MilliQ water. The glass was blown dry using compressed air and then incubated overnight at 50°C. Sample chambers were formed by creating narrow channels with double-sided tape with approximate volumes of 5 µL. RNA samples were diluted in buffer in a similar manner as described for annealing experiments. The RNA solution was then carefully flowed into the channel by pipetting. Oil was used to seal either side of the channel to prevent evaporation. Samples were immediately imaged using temperature-controlled microscopy.

#### Temperature-controlled microscopy measurements

Samples were placed inside a custom temperature-controlled stage (Instec) with temperature ranging from 0°C to 90°C. The temperature was measured using a K-type thermocouple with a

USB digital to analog converter (DAQ) designed for use with a thermocouple (Measurement Computing). The DAQ software creates CSV files with four temperature points per second. The temperature of the sample was measured using a custom 3-D printed jig to ensure probe placement on the top glass surface near the sample. The effective temperature range at the sample location in a 20 min experiment was 10°C to 80°C with temperatures between 0°C and 85°C taking an additional 40 min to achieve. Videos were acquired on a Zeiss Primovert microscope set to brightfield imaging using a 40X air objective. Micromanager was used to capture images at one frame per second with a Blackfly S camera (Teledyne FLIR) mounted to a custom 3D printed 1X magnification adapter. Alternatively, a Zeiss Primovert was used with a 40X air objective and a Zeiss Axiocam camera with a 0.6X adapter. Temperature files were aligned to images by comparing the start time and date in the image file metadata with the timestamp in the CSV. Temperatures, where phase transitions were observed (i.e., $T_{phase}$), were also manually recorded during acquisition. The reported $T_{phase}$ was determined by at least three technical replicates.

#### Fluorescent recovery after photobleaching experiments

Samples were prepared by either temperature-controlled microscopy or by annealing in a chamber prepared as described above. Droplets were allowed to settle onto the coverglass surface for 5 min and then the sample was placed on a confocal microscope (C-trap, Lumicks) and imaged with a 63X oil objective. Regions for continuous imaging, a 4-frame reference, and bleaching were selected using Bluelake software (Lumicks). Samples were bleached for 8 frames with a 150 ms dwell time per pixel. Recovery was observed for 5 min post-bleach. Analysis of FRAP images was performed via python using the numpy and scipy libraries. Normalization and correction of intensities were performed as described by Taylor et al[10].

#### Simulations

##### Force field parameters for polyphosphate, $Mg^{2+}$ and $Cl^-$ ions

The chemical structure of polyphosphate used in simulations, $PO_3^{2-}$-$(PO_3^-)_{16}$-O-$PO_3^{2-}$, is shown in **Supplementary Fig. S4a**. The bonded interaction parameters were derived by ACPYPE [11], and amber2 was chosen for the "atom_type" option. The atom types are shown in **Supplementary Fig. S4b**. The partial charges of each atom were fit using an online R.E.D. Server [12], which allows users to perform quantum calculations of small molecules, including geometry optimization and single-point energy calculations. The RESP method[13] was implemented to derive partial charges. The molecule $PO_3^{2-}$-$PO_3^-$-O-$PO_3^{2-}$ (**Supplementary Fig. S4c**) was used to fit partial charges for the atom types O2, OS and P. The partial charges for atom types at two termini ($O2^T$ or $P^T$) are different from those in the middle part ($O2^M$ and $P^M$). Specifically, $O2^T$ and $O2^M$ share the same force field parameters except for their partial charges. Likewise, $P^T$ and $P^M$ share the same force field parameters except for their partial charges. RESP-B1 was chosen for the keyword "CHR_TYP" in the input file of the job submitted to the R.E.D. Server. This approach mirrors that described by Duan et al. [14] to fit the partial charges. First, the geometry of $PO_3^{2-}$-$PO_3^-$-O-$PO_3^{2-}$ was optimized at the HF/6-31G** level of quantum calculations. Then, B3LYP exchange and correlation functionals [15,16] with the ccpVTZ basis set [17] were used for the single-point energy calculations. Finally, RESP method[13] was used to derive the partial charges. The total charge of $PO_3^{2-}$-$PO_3^-$-O-$PO_3^{2-}$ was −5.0. A charge constraint was applied so that the sum of partial charges in the middle phosphate group is −1.0, namely $q(OS)+q(P^M)+2*q(O2^M) = -1.0$, where q denotes the value of partial charges. The final fitting results are listed in **Supplementary Table S2**. The derived partial charges of $P^M$ and $O2^M$ were used for P and O2 atoms in the repeating unit of polyphosphate, and partial charges of $P^M$ and $O2^M$ were used for P and O2 atoms in the two termini of polyphosphate. The van der Waals radii of O2 and OS atoms were taken from Steinbrecher et al. [18]. It has been shown before that these radii of phosphate oxygen atoms can better describe RNA structures and avoid spurious hydrogen bonds[19]. The force fields of $Mg^{2+}$ and $Cl^{-}$ were taken from Grotz et al.[20]. The "*microMg*" parameters[20] were used for $Mg^{2+}$ for simulations with TIP3P water model [21]. This set of parameters can reproduce a wide range of experimental data, including the solvation free energy, the distances to the oxygens of the first hydration shell, the hydration number, the activity derivative in $MgCl_2$ solutions and the self-diffusion coefficient [20].

**Details of the setup of molecular dynamics simulations**

All simulations were performed using the Gromacs 2021 package [22, 23, 24] on the GPU nodes of RIS cluster at Washington University in St. Louis. A linear structure of $PO_3^{2-}$-$(PO_3^{-})_{16}$-O-$PO_3^{2-}$ was first solvated in a cubic water box with the size of $7 \times 7 \times 7\ nm^3$. Then, 10 $Mg^{2+}$ counterions and additional 100 mM $MgCl_2$ were added to the system. The TIP3P water model [21] was used for the simulation. The force field parameters for polyphosphate and ions are described in the section above. The overall energy of the system was minimized using the steepest descent method followed by 500 ps NPT equilibration simulations at 298 K and 1 bar. The equilibrated structure was used as the starting structure for multiple independent NPT simulations with different initial velocities. Ten independent NPT simulations were performed for each temperature and each independent simulation was 500 ns long. The V-rescale thermostat [25] was used for maintaining the temperature with a coupling time constant of 0.1 ps. The Parrinello-Rahman method [26, 27] was used to maintain the pressure at 1 bar with a coupling time constant of 2.0 ps. Periodic boundary conditions were used and PME algorithm [28, 29] was applied to calculate the long-range electrostatic interactions. The cutoffs of the short-range electrostatic potential and van der Waals potential were set to 1.2 nm and 1.1 nm, respectively. The LINCS algorithm [30] was used to constrain all bonds with H-atoms.

#### Supplementary Figures

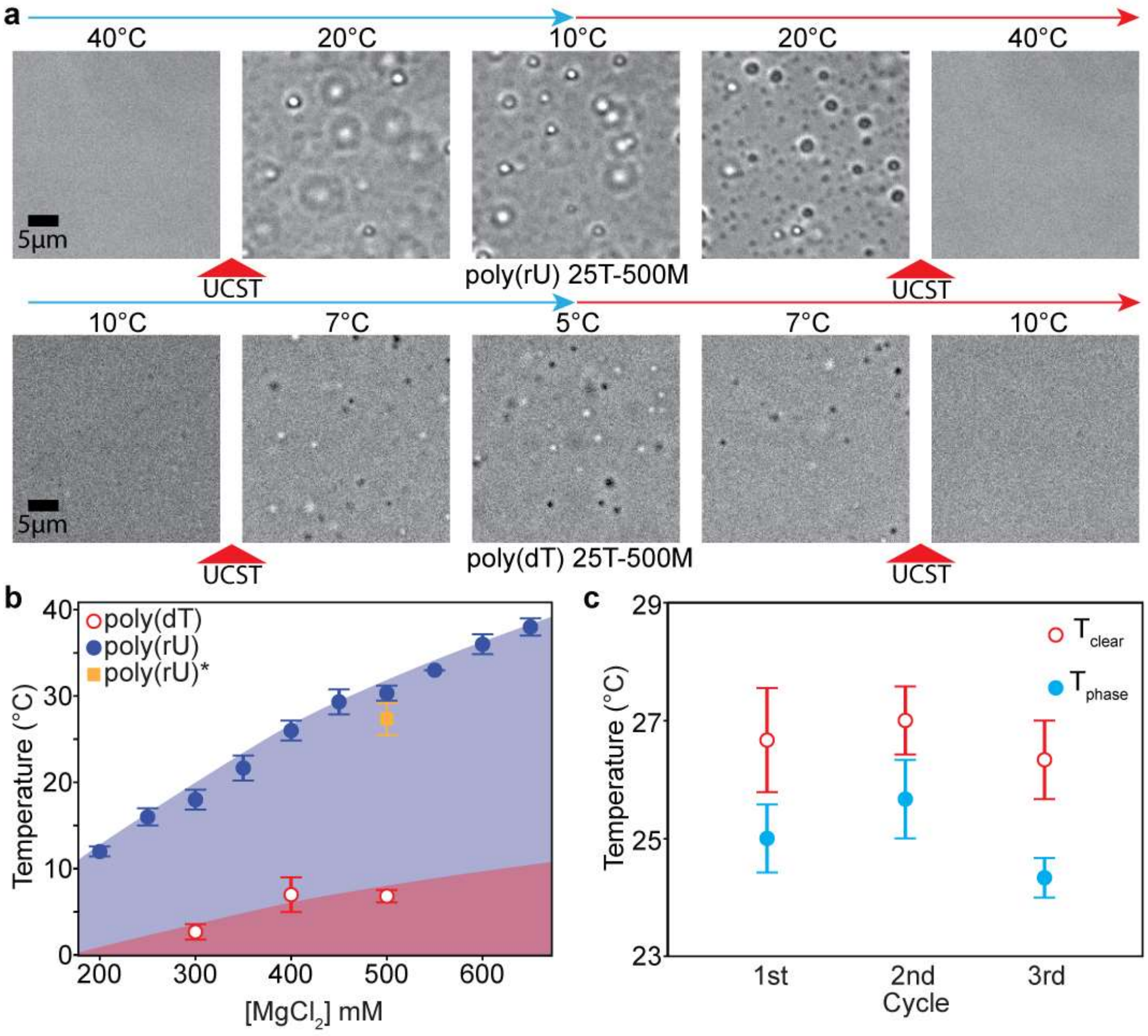


**Supplementary Figure S1. a.** A comparison of 1.5 mg/mL poly(rU) to poly(dT) in 25 mM Tris-HCl (pH 7.5 at 25°C), 500 mM $Mg^{2+}$. Poly(dT) solution undergoes a reversible UCST-type transition at 6.8 ± 0.7°C. **b.** A state diagram of poly(rU) RNA (1.5 mg/mL) at varying concentrations of $Mg^{2+}$ with 25 mM Tris-HCl (pH 7.5 at 25°C). Data (blue, filled circle) replotted from Pullara et al[31]. The shaded region indicates where phase separation occurs. The yellow square data point is derived from Movie 1. **c.** Sensitivity of the $T_{phase}$ and $T_{clear}$ of poly(rU) to repeated phase transitions. Samples of poly(rU) were oscillated around the $T_{phase}$ and showed no substantial change in the $T_{phase}$ over 3 complete cycles. Buffer notation used: the number in front of "T" indicates [Tris-HCl] and the number in front of "M" indicates [$Mg^{2+}$] in a given buffer.

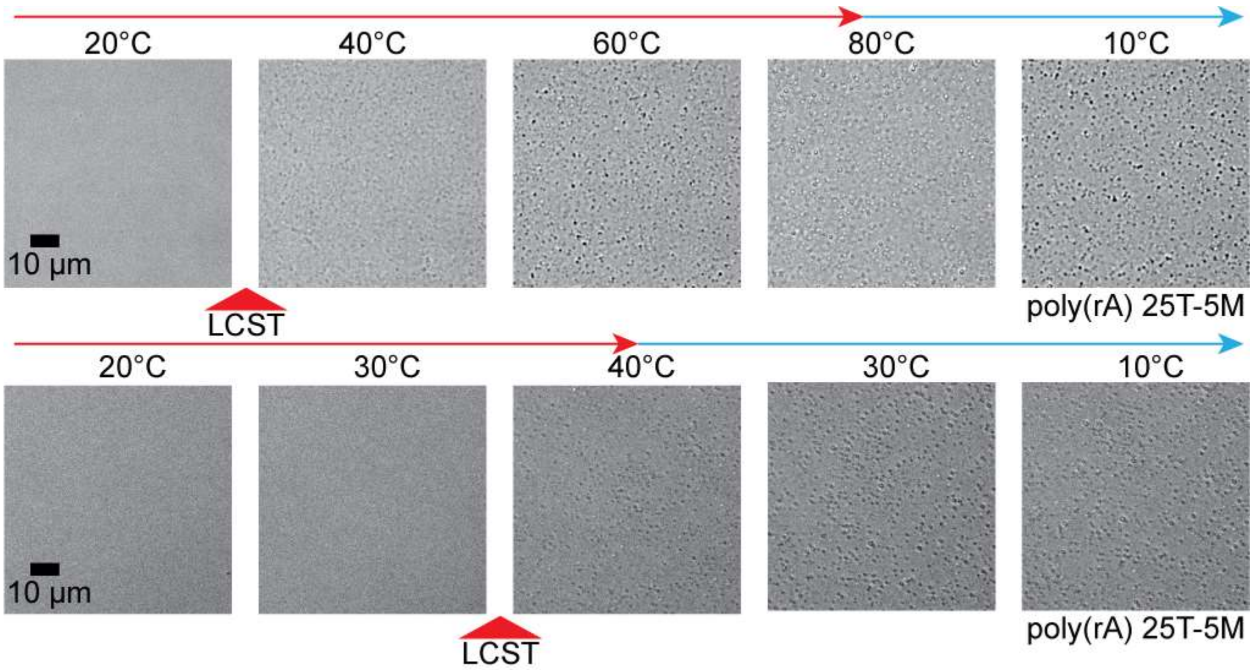


**Supplementary Figure S2. Phase separation and dynamical arrest of poly(rA) condensates**. The images shown here correspond to 1.5 mg/mL poly(rA) in 25 mM Tris-HCl (pH 7.5 at 25°C) and 5 mM $Mg^{2+}$, undergoing a temperature increase from 20°C to 80°C (top) or 45°C (bottom) and a subsequent cooling to 10°C. Note that droplets persisted even after cooling below the cloud point temperature (39.3 ± 8.5°C).

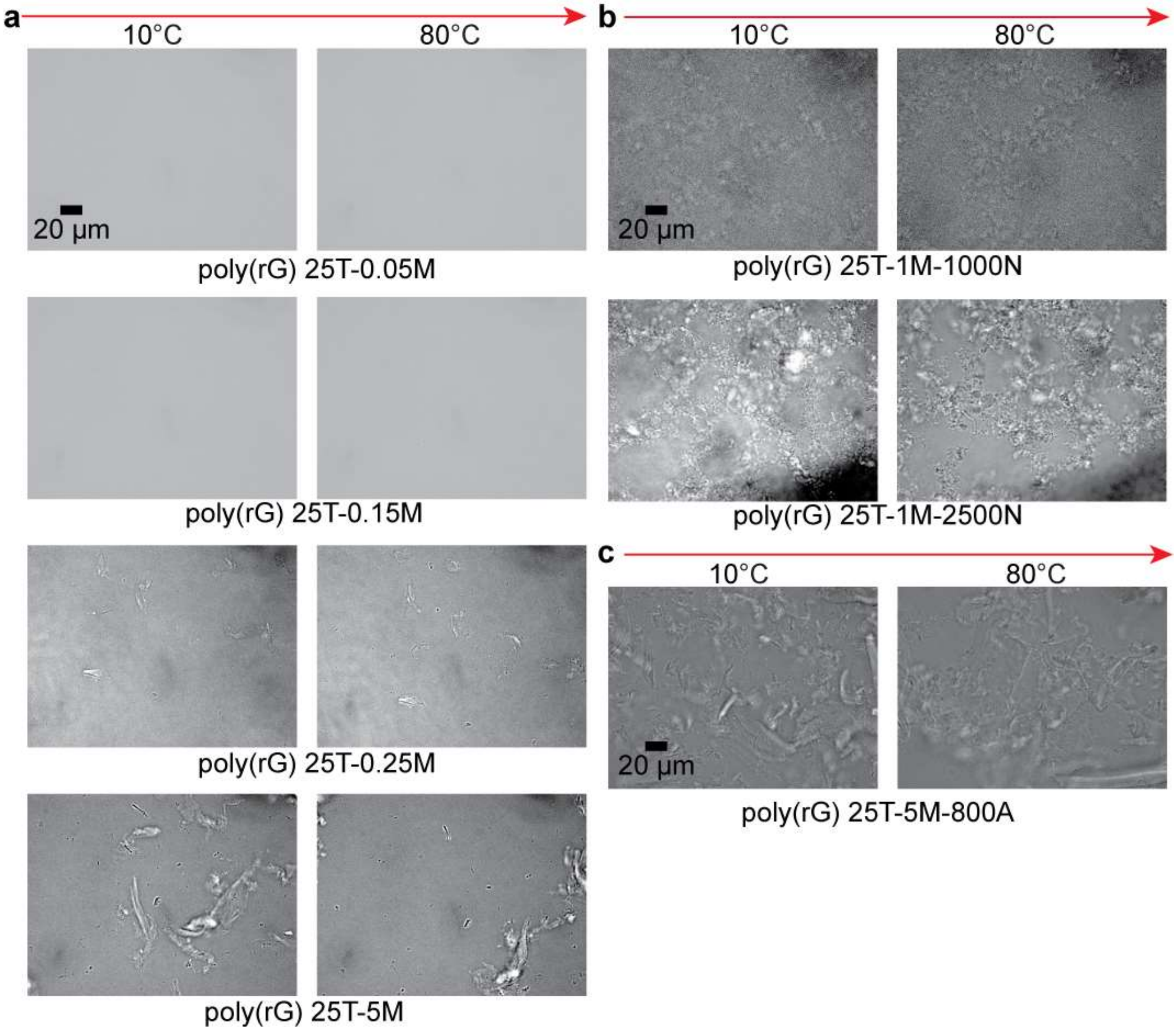


**Supplementary Figure S3. $Mg^{2+}$ sensitivity of poly(rG) aggregates**. **a.** Samples of 1.5 mg/mL poly(rG) in 25 mM Tris-HCl (pH 7.5 at 25°C) were subjected to a temperature ramp from 5°C to 80°C. At ≤ 0.15 mM $Mg^{2+}$, we observed no aggregation. **b.** Upon addition of NaCl, we observed some variation in the morphology of the poly(rG) aggregates. Additionally, we observed changes in the network structure of the aggregates as the temperature was increased. **c.** With the addition of ammonium acetate, we observed no significant variation in poly(rG) aggregation. Buffer notation used: the number in front of "T" indicates [Tris-HCl], the number in front of "M" indicates [$Mg^{2+}$], the number in front of "N" indicates [$Na^+$], and the number in front of "A" indicates [AmAc] in a given buffer.

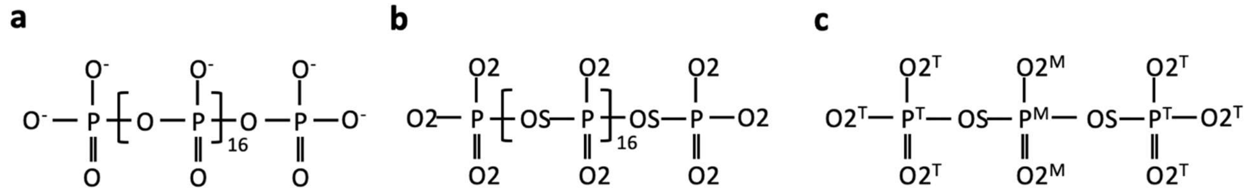


**Supplementary Figure S4. a.** Chemical structure of polyphosphate [$PO_3^{2-}$–$(PO_3^{-})_{16}$-O-$PO_3^{2-}$] used in simulations, and **b.** corresponding atom types. **c.** The molecule used to fit partial charges. Please note that the partial charges for atom types at two termini ($O2^T$ and $P^T$) are different from those in the middle region ($O2^M$ and $P^M$).

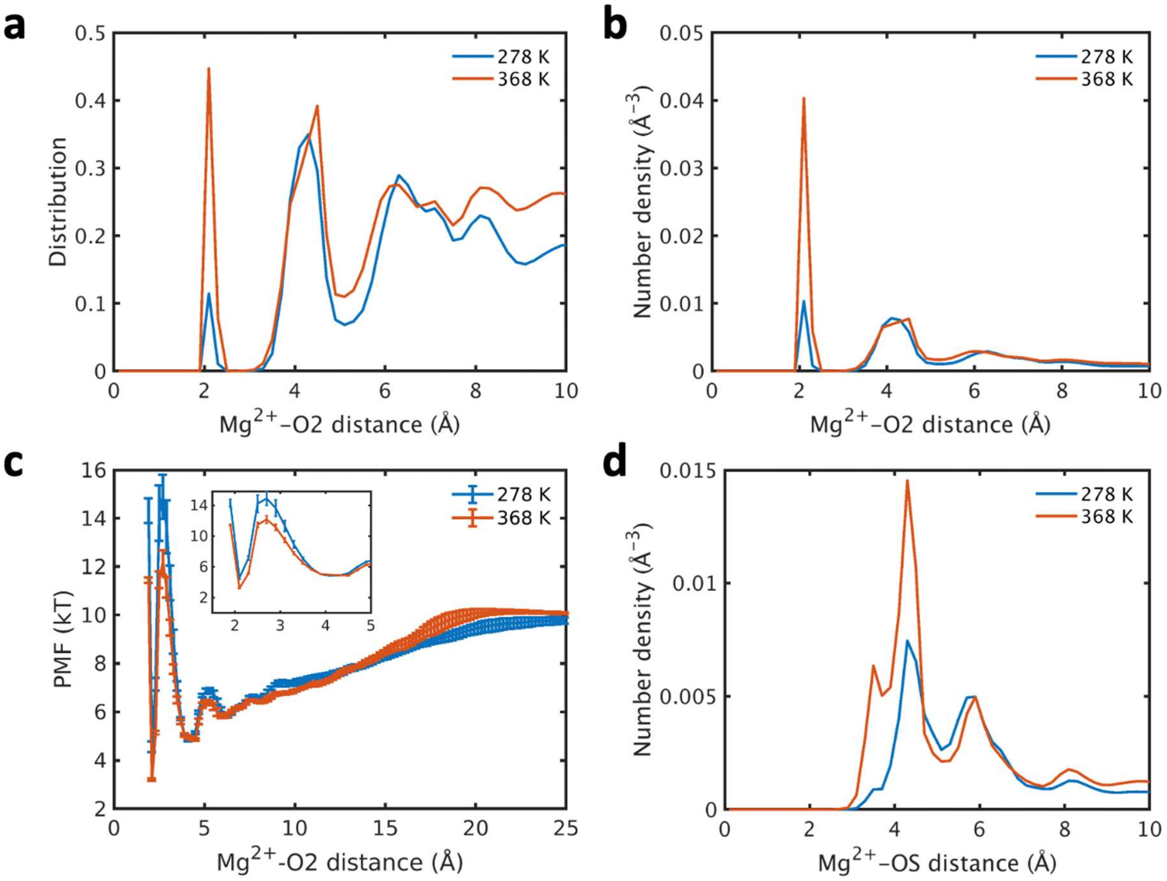


**Supplementary Figure S5. More $Mg^{2+}$ ions bind to O2 atoms of polyphosphate at a higher temperature. a.** Ensemble-averaged distribution of $Mg^{2+}$ ions around the O2 atom *P(r)* as a function of the distance between the $Mg^{2+}$ ion and O2 atom. *P(r)* is calculated as the histogram of all the pairwise distances between $Mg^{2+}$ ions and O2 atoms in the system, which is further averaged by the number of O2 atoms. The bin width for the histogram is 0.2 Å. **b.** Distribution of number density of $Mg^{2+}$ ions around the O2 atom *n(r)*, $n(r) = \frac{P(r)}{4\pi r^2 dr}$, where *r* is the distance between the $Mg^{2+}$ ion and O2 atom and *dr* is the bin width, 0.2 Å. **c.** Potential of mean force (PMF) between the $Mg^{2+}$ ion and O2 atom, which is calculated as $-kTlog\big(n(r)\big)$. The inset zooms in the region from *r = 1.5 Å* to *r = 5 Å*. Errors reported reflect the standard deviation of the results from ten independent simulation trajectories. **d.** Ensemble-averaged distribution of $Mg^{2+}$ ions around the OS atom as a function of the distance between the $Mg^{2+}$ ion and OS atom. The calculation of this distribution is same as **b**.

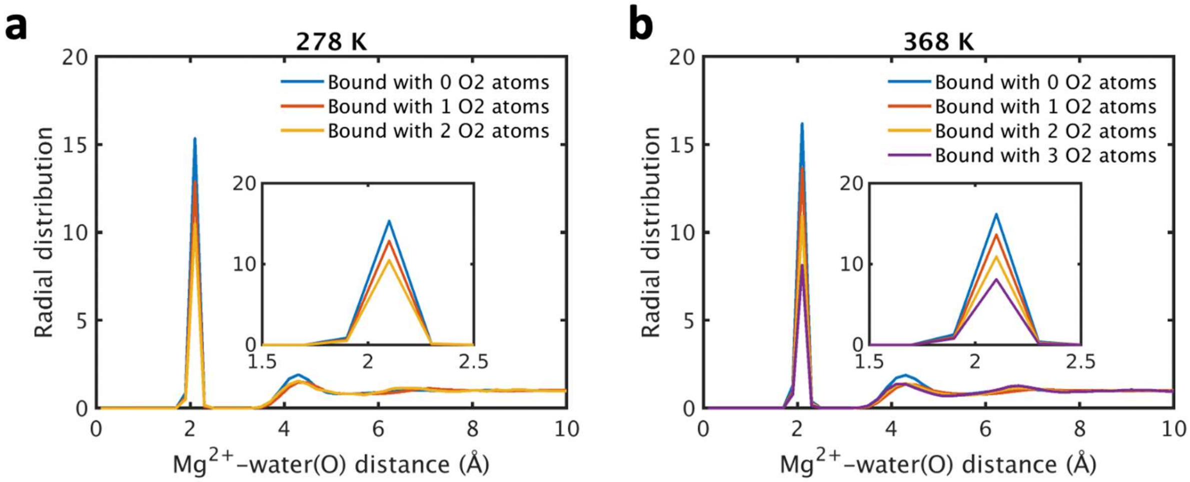


**Supplementary Figure S6. Binding of O2 atoms to a $Mg^{2+}$ ion releases waters in the first solvation shell of the $Mg^{2}$ ion.** Radial distribution function of oxygen atoms (water(O)) around $Mg^{2}$ ions with different coordination numbers at **a.** 278 K and **b.** 368 K. The insets zoom in the region from 1.5 Å to 2.5 Å. These results show fewer waters in the solvation shell of the $Mg^{2+}$ ion when more phosphate groups bind to the $Mg^{2+}$ ion.

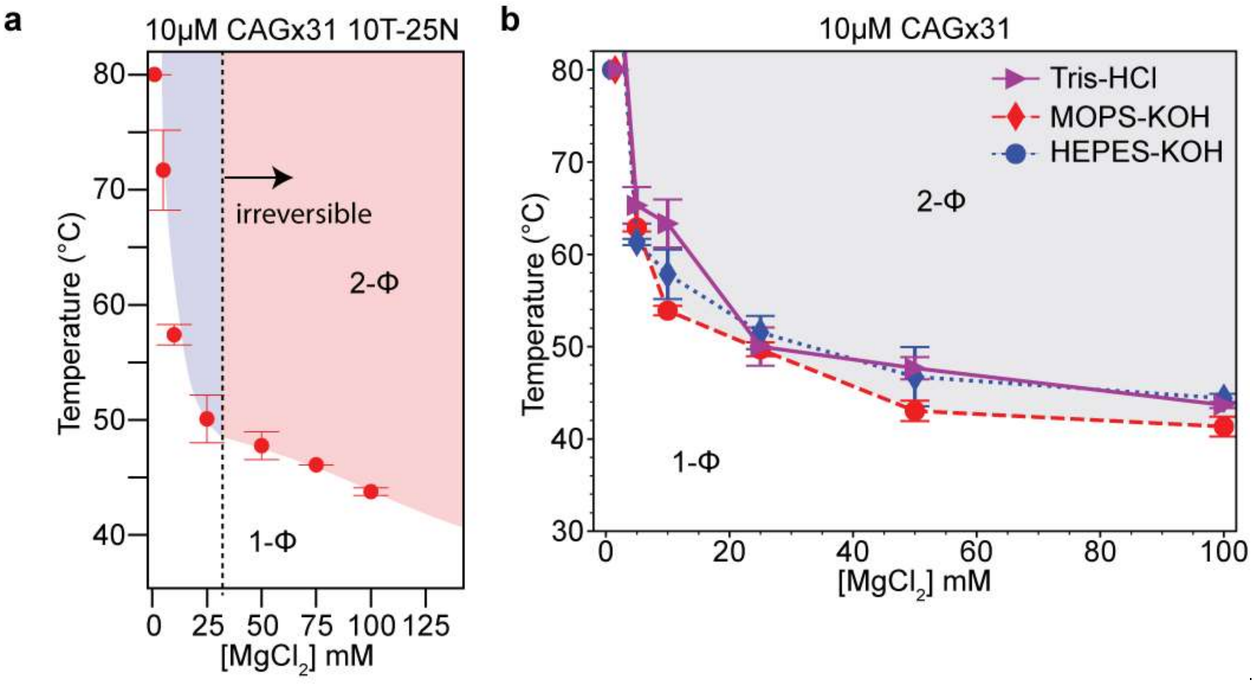


**Supplementary Figure S7.** State diagrams for CAGx31 at variable $Mg^{2+}$ and buffering agents. **a.** A state diagram for CAGx31 (inset to **Fig. 3e** in the maintext). The cloud point temperatures are plotted as a function of [$Mg^{2+}$]. Conditions which induced irreversible and reversible phase separation are shown with red and blue shading, respectively. **b.** A comparison of Tris-HCl as a buffering agent to alternative buffers which display less pH variation in response to temperature. Each titration of [$Mg^{2+}$] was carried out in 25mM of either Tris-HCl (pH 7.5 at 25°C) (*purple triangle, solid line*), MOPS-KOH (pH 7.5 at 25°C) (*red diamond, dashed line*), or HEPES-KOH (pH 7.5 at 25°C) (*blue circle, dotted line*). The gray shaded region is a guide to the eye representing the two-phase regime. Buffer notation used: the number in front of "T" indicates [Tris-HCl] and the number in front of "N" indicates [$Na^+$] in the buffer.

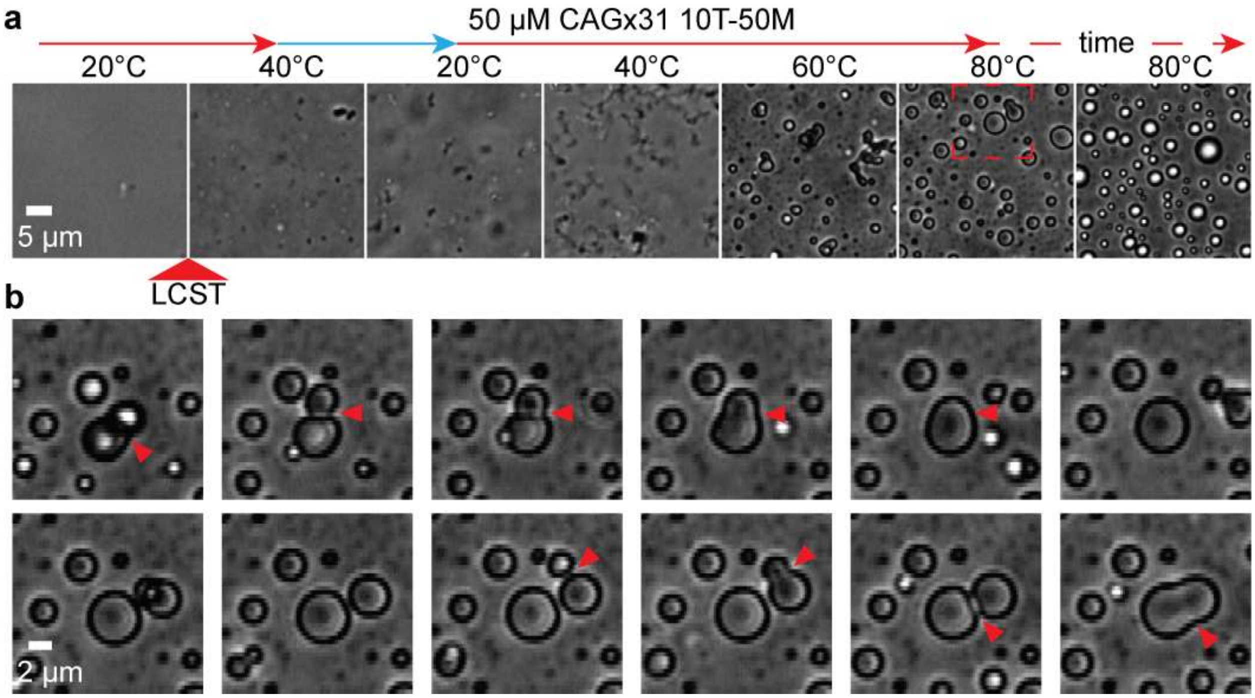


**Supplementary Figure S8**. **Liquid-like properties of CAGx31 droplets at elevated temperatures**. **A.** Fifty µM CAGX31 phase separated at 41.1 ± 1.7°C in a buffer containing 10 mM Tris-HCl (pH 7.5 at 25°C) and 50 mM $Mg^{2+}$ forming aspherical condensates that persisted even when the temperature was lowered immediately below its cloud point temperature. This sample was then heated again. At elevated temperatures, aspherical condensates relaxed into spherical droplets. **b.** Liquid-like properties of RNA droplets. A zoomed in view of several droplets from the 50 µM CAGX31 shown in **a (red box)**. When incubated at 80°C, droplets demonstrated rapid fusion. Droplets merged and relaxed repeatedly over several minutes. The selected time-lapse images are extracted from Movie 9.

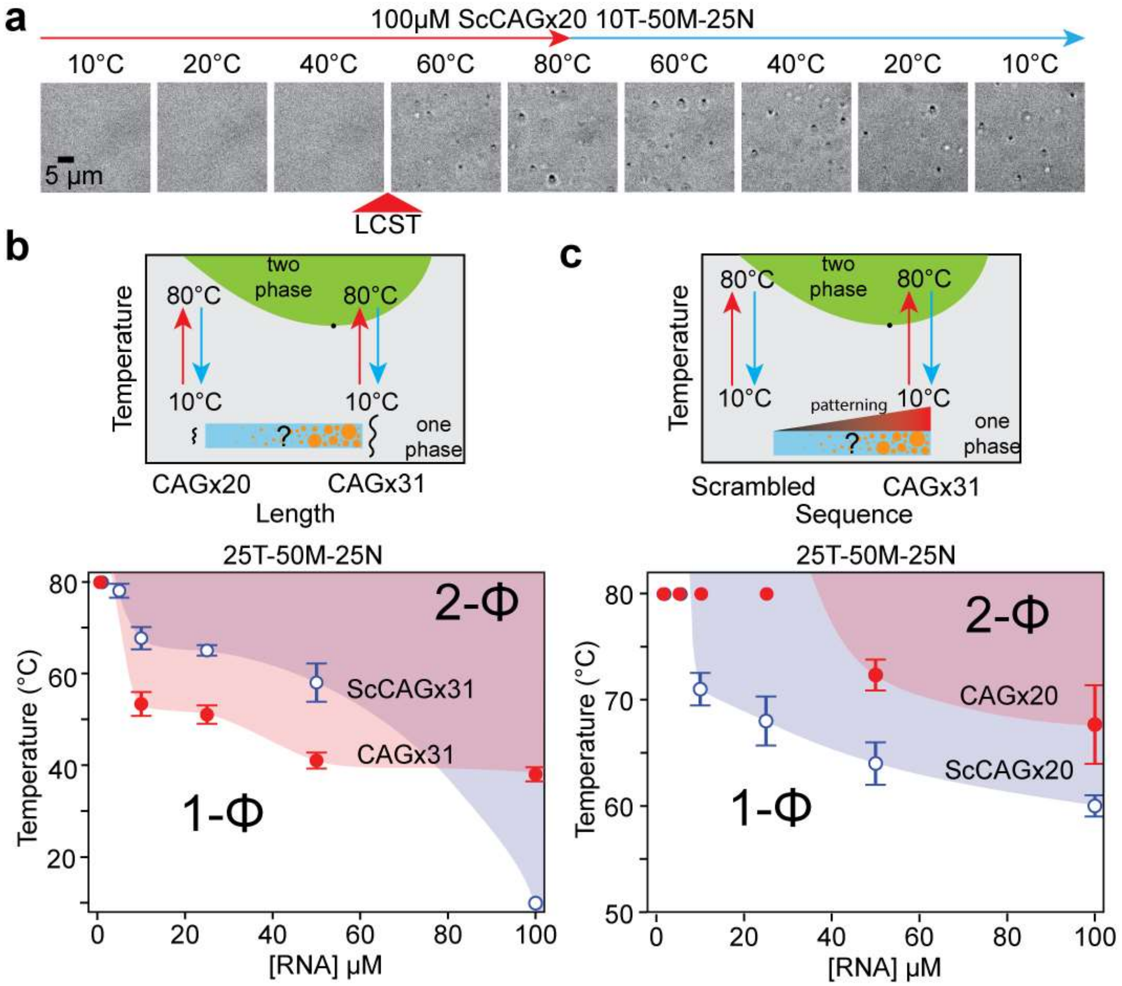


**Supplementary Figure S9. a**. Irreversible phase separation of a scrambled RNA containing CAG equivalent to CAGx20 (See Supplementary Table S1). The scrambled CAGx20 sequence underwent phase separation at 60.1 ± 2.2°C in a buffer containing 25 mM Tris-HCl (pH 7.5 at 25°C), 50 mM $Mg^{2+}$, and 25 mM $Na^{+}$. Droplets did not dissolve upon cooling below the LCPT of the sample. **b.** Length and sequence dependence of LCPT of CAG repeat RNA. A state diagram showing the LCPT as a function of RNA concentration for CAGx31 and CAGx20 (colored red, left and right panels) versus a scrambled sequence of equivalent CAG content to CAGx31 or CAGx20 (colored blue, left and right panels, respectively). Buffer notation used: the number in front of "T" indicates [Tris-HCl], the number in front of "M" indicates [$Mg^{2+}$], and the number in front of "N" indicates [$Na^{+}$] in a given buffer.

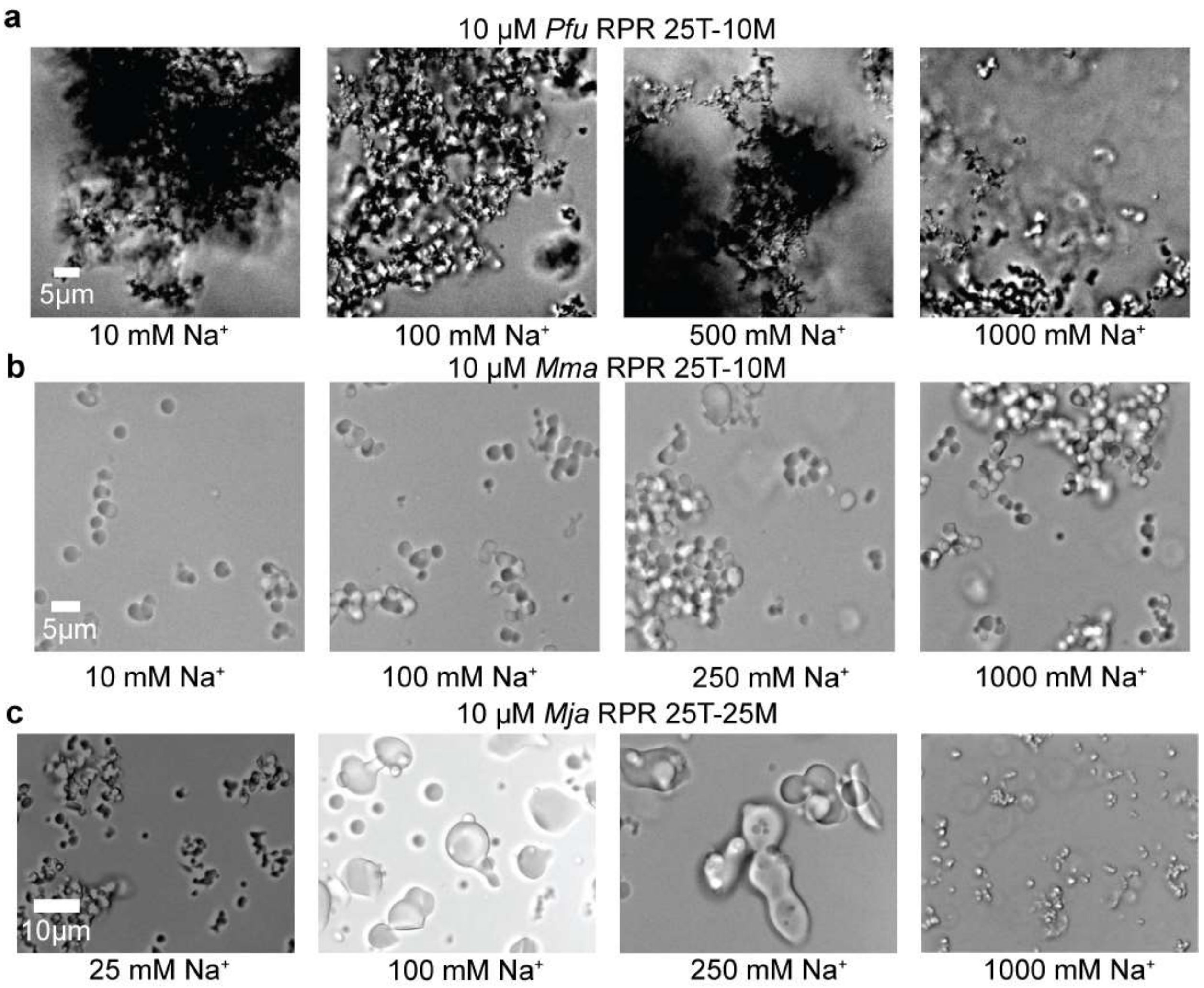


**Supplementary Figure S10**. **Effect of $Na^+$ on RPR condensate morphology**. **a**. *Pfu* RPR; **b**. *Mma* RPR; **c**. *Mja* RPR. While *Mja* RPR was prepared in a buffer containing 10 mM Tris-HCl (pH 7.5 at 25°C) and 25 mM $Mg^{2+}$, *Mma* and *Pfu* RPRs were in 10 mM $Mg^{2+}$. For *Mja* RPRs, few condensates were observed at 10 mM $Na^+$. In contrast to the morphology of *Mja* RPR droplets that underwent visible changes upon varying [$Na^+$], there was little change in *Mma* or *Pfu* RPR droplets upon increasing [$Na^+$]. Buffer notation used: the number in front of "T" indicates [Tris-HCl] and the number in front of "M" indicates [$Mg^{2+}$] in a given buffer.

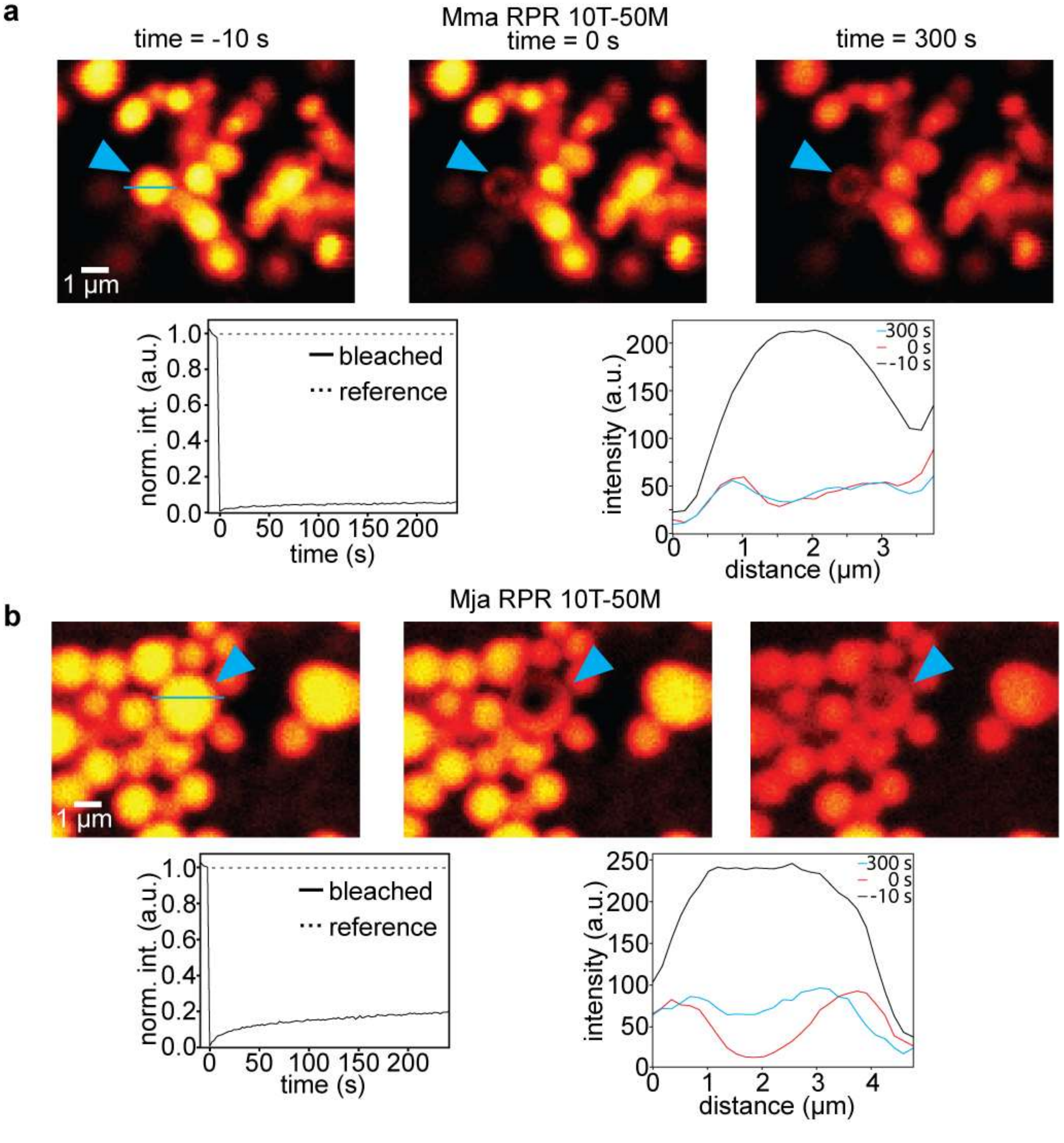


**Supplementary Figure S11.** Fluorescent images to show fluorescence recovery after photobleaching (FRAP) of *Mma* and *Mja* RPRs. **a.** *Mma* RPR; **b.** *Mja* RPR. Ten µM fluorescein-labeled *Mma*/*Mja* RPR was subjected to FRAP in a buffer containing 10 mM Tris-HCl (pH 7.5 at 25°C) and 50 mM $Mg^{2+}$. Images were acquired over 300 s. While no significant fluorescence recovery was observed with *Mma*, there was ~20% recovery compared to the reference in the case of *Mja* RPR.

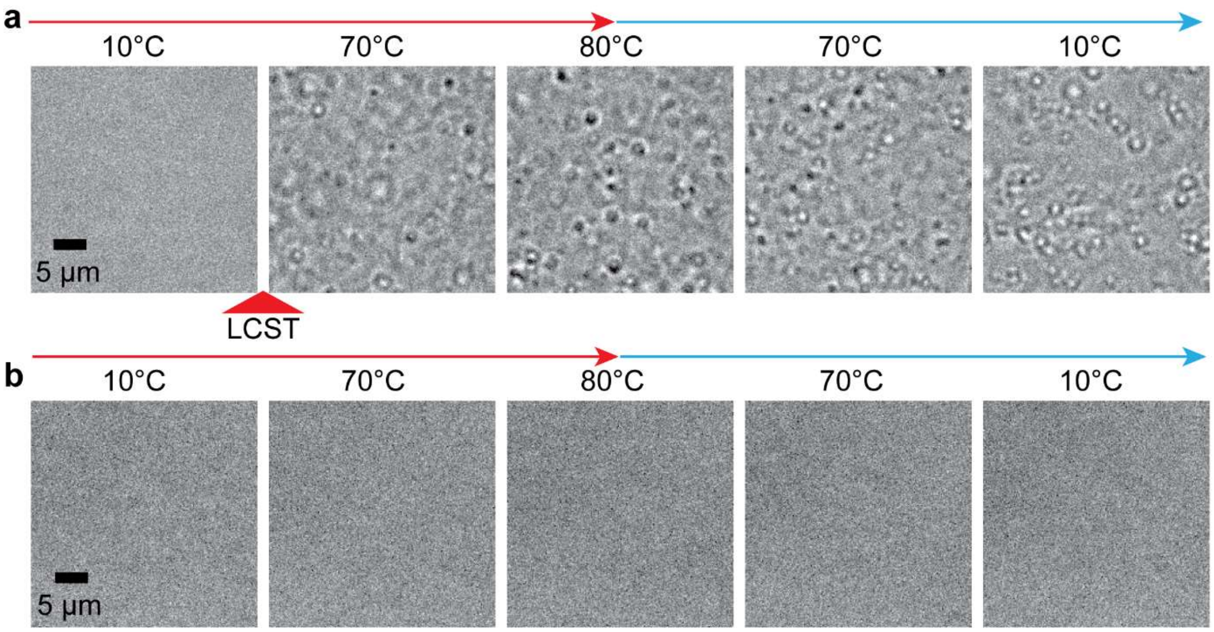


**Supplementary Figure S12. Buffer dependence of *Pfu* RPR phase separation**. **a.** *Pfu* RPRs (10 μM) underwent irreversible phase separation in a buffer containing 50 mM HEPES-KOH (pH 7.5 at 25°C) and 50 mM $Mg^{2+}$. Phase separation was observed at 68.2 ± 1.3°C and droplets did not dissolve upon cooling below the LCPT of the sample. **b.** *Pfu* RPR was imaged in a buffer containing 50 mM HEPES-KOH (pH 7.5 at 25°C), 50 mM $Mg^{2+}$, and 0.8 M $NH_4OAc$. Phase separation did not occur during one round of heating and cooling, suggesting that ammonium acetate suppresses *Pfu* RPR phase separation.

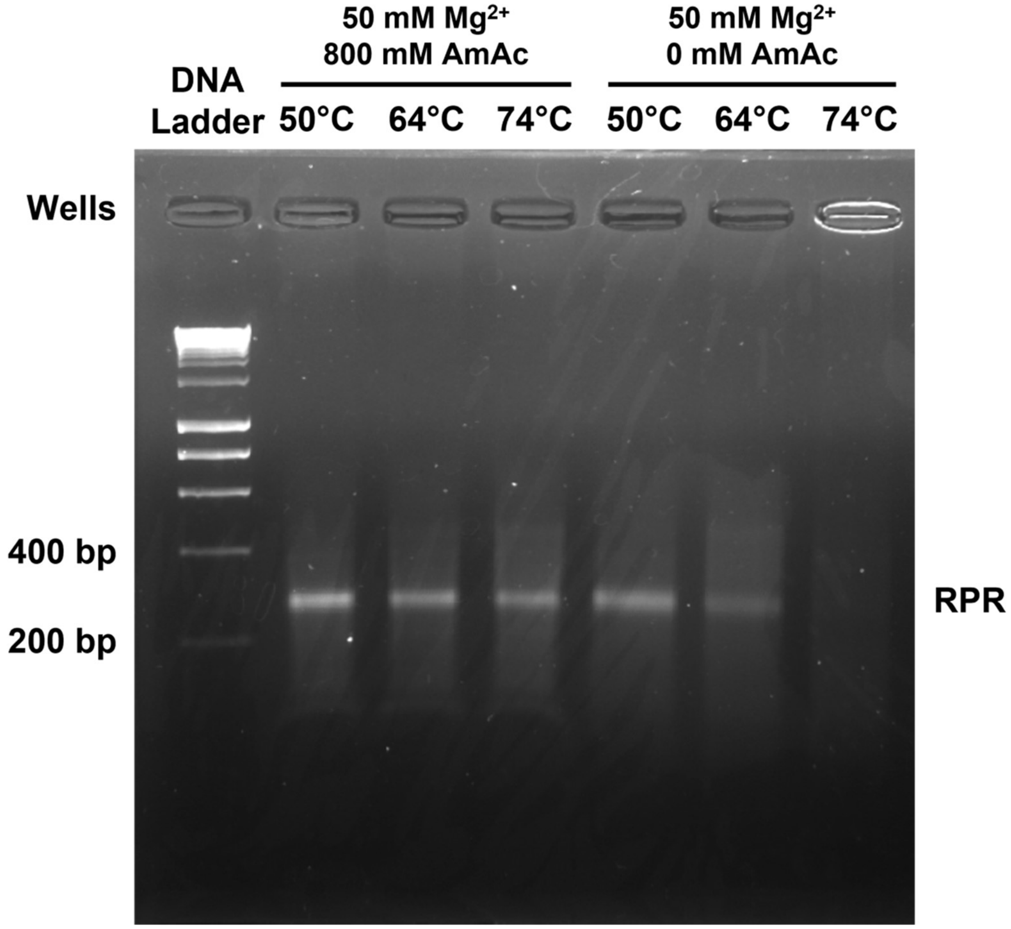


**Supplementary Figure S13.** Gel image of *Pfu* RPR after annealing under conditions identical to those used in pre-tRNA cleavage assays (**Fig. 6b** in the maintext). *Pfu* RPR was annealed in 50 mM HEPES-KOH (pH 7.5 at 55°C) and 50 mM $Mg^{2+}$ at temperatures bracketing $T_{phase}$ (69°C) as determined by microscopy experiments. Each assay was performed in the presence or absence of 800 mM ammonium acetate (AmAc), which suppresses phase separation at the $T_{phase}$ +5°C temperature. In the absence of AmAc where phase separation is observed, the band intensity diminishes with increased temperatures approaching or exceeding the $T_{phase}$. Notably, at $T_{phase}$ +5°C (74°C), the presence of a high molecular weight RNA trapped in the loading well is consistent with the observation of micron-sized percolated RNA condensates observed in microscopy experiments (**Fig. 5** in the maintext; **Supplementary Fig. S12a**).

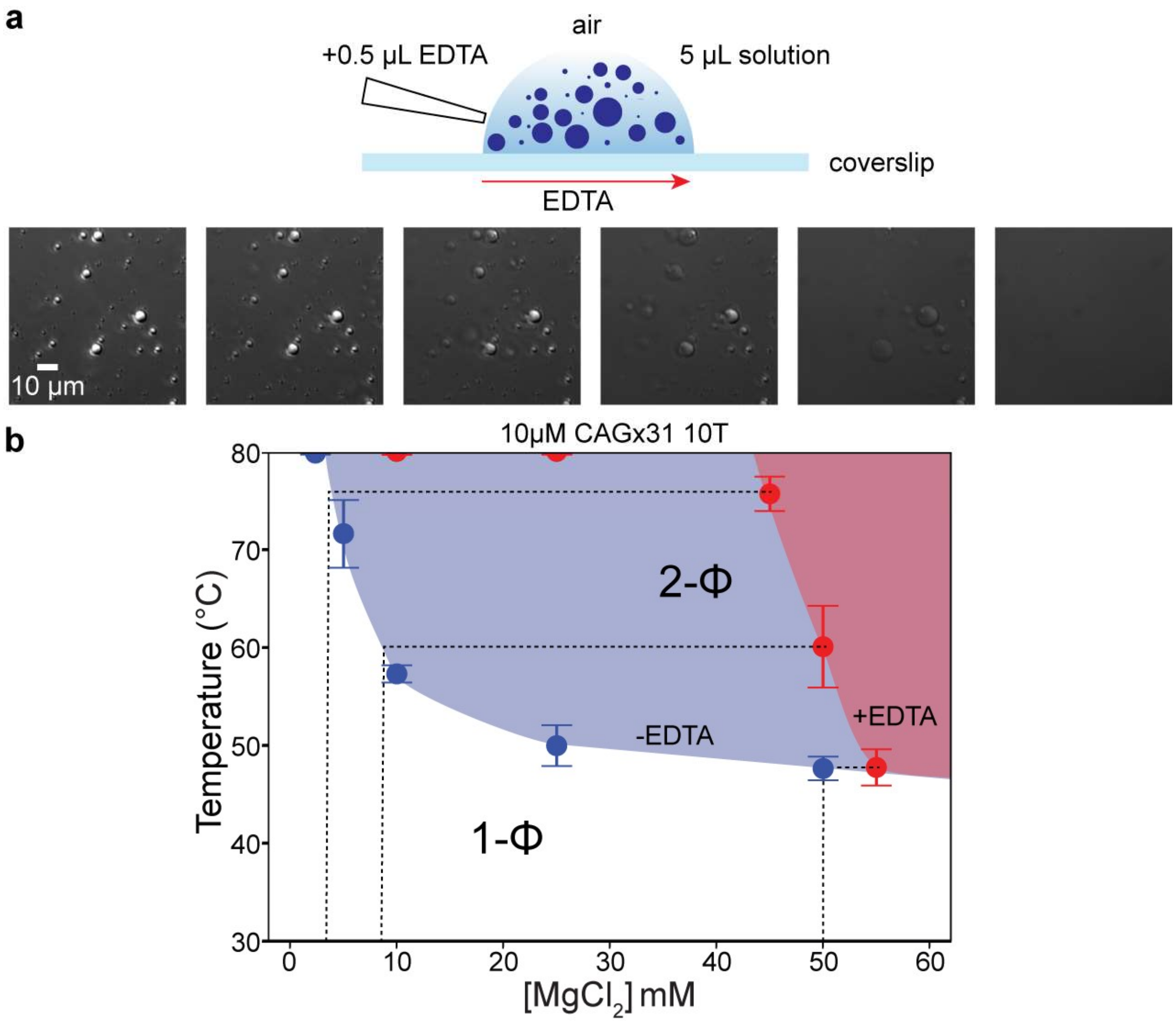


**Supplementary Figure S14. Effect of EDTA on phase separation and percolation of CAGx31 RNA. a.** EDTA-mediated dissolution of arrested RNA droplets. Shown here are DIC images of a solution of 10 µM CAGx31 RNA that underwent annealing in a buffer containing 10 mM Tris-HCl (pH 7.5 at 25°C) and 50 mM $Mg^{2+}$. Subsequently, a small volume of 500 mM EDTA was added to an open-air droplet on a coverslip and the diffusion of EDTA through the solution was immediately observed on a microscope (*see* Movie 25). The volume of EDTA was chosen to achieve a 1:1 ratio of EDTA:$Mg^{2+}$. The RNA droplets dissolved in the image frame from left to right corresponding to the direction of EDTA diffusion relative to the imaging area. **b.** EDTA modulates CAGx31 thermoresponsive phase separation. The same sample conditions as in **a** were compared with the addition of 50mM EDTA.

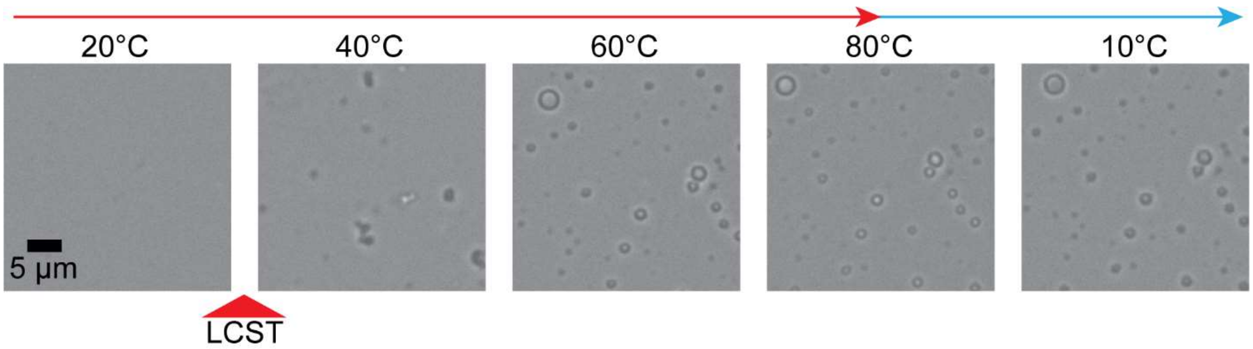


**Supplementary Figure S15.** Phase separation of CAGx31 RNA in the presence of dimethylsulfoxide (DMSO). Shown here is a solution of 10 µM CAGx31 RNA that underwent one cycle of heating and cooling in a buffer containing 10 mM Tris-HCl (pH 7.5 at 25°C) and 50 mM $Mg^{2+}$, with the addition of 5% (v/v) DMSO (*see* Movie 26). The cloud point temperature of the sample shifted substantially from 47.7 ± 1.2°C (without DMSO) to 29.4 ± 3.3°C (with DMSO).

**Supplementary Table S1**

| Name | Sequence (5' to 3') | Source |
|---|---|---|
| *Pyrococcus furiosus* (*Pfu*) RNase P RNA (RPR) | ggcaaaucaaccuaaaacuuacacaGGCGAGGGGGCUGGGGGCUGUCGG<br>GCUCGUGCCCGAGGAAGUUCCGCCCACCGCACCGGGGCCGCGGUGCCGUAA<br>GGCACCGGCCGAGAGGCCGGGCAACGGCACAGAAACGACACGUCCCUCGGG<br>GGAUGUGGAUGAAAGCGGUGAAGGCUCCCGGUGACGGGAGCCGAGUUAACC<br>CGCAGACAAUCCCGAGGGGAGCGGUGAAACGGCCGUCCCGCGGGGUGCAAG<br>GCCGAGUUAGGGCCGAUGAGUUCCCGGUGUGAGGCCCGUGGUAGGCCGCU<br>UAGUCGAAUGCUCCCGUAGUACAGAAGGCGGGCUAUAGCCCCCUCGCCGGA<br>GUAUGGUAAACUUUUUGUUCGAAUU | IVT |
| *Methano-coccus maripaludis* (*Mma*) RPR | ggcaaaucaaccuaaaacuuacacaUCGUGGCAAGGGGGCUGGUGACUA<br>UCCACUGGGAGGAAGCUCUGCCCACCCAAUUGUGUAGAUUUCCUGAGAAGG<br>AAUGAAUAGGGUAUAGAAACGACACGGUUCUAAGAAAUAUGACUAUGAUAUU<br>UUCAAAAUUGAGGAUUCUUAGAAAAUCGAUGAAACGACCUUAUUUUACACGG<br>GUGCAAGCAUAUGCGCUAAGCUGAAUGUCACCGAAAUUACAGAAGGCAGGCU<br>AUAGCCCCGAUGCCAUUUCGAGUAUGGUAAACUUUUUGUUCG | IVT |
| *Methano-caldo-coccus jannaschii* (*Mja*) RPR | ggcaaaucaaccuaaaacuuacacaGGCGGGUAAGGGGGCUGGUGAC<br>UUUCCCCUCUUUAAGAGGGGAGGAAGUUCCGCCCACCCCAUUUAUGGGCAG<br>CGUCCCCUGAGAAGGGGCGGGAGAUGCAGCAGAAACGACACGGCUCCGGAA<br>GAGAUGACGAUGAUAGUGAAAGUUGAGGACUUCCGGAGAACCGGUGAAACG<br>GGCAUCUCCCCUGCCCGGGGUGCAAGCCGGUUUCGGCGCUUAGCCGAAUGU<br>CACCGAAAUUACAGAAGGCGGGCUAUAGCCCCCAUACCCGCGAGUAUGGUAAACUUUUUGUUCG | IVT |
| CAG 20 | CAGCAGCAGCAGCAGCAGCAGCAGCAGCAGCAGCAGCAGCAGCAGCAGCAGCAGCAGCAG | IVT |
| Scrambled CAG 20 | GGCGAGGAACGACACAGGACCAAAAAGAAAGCGGCCACCGCCGGACACGCGCACGCAGCGGG | IVT |
| CAG 31 | CAGCAGCAGCAGCAGCAGCAGCAGCAGCAGCAGCAGCAGCAGCAGCAGCAG<br>CAGCAGCAGCAGCAGCAGCAGCAGCAGCAGCAGCAGCAGCAG | IVT |
| Scrambled CAG 31 | AACACGCGCCCGAACCCAAAGGCAACCCAGGAACGAAAAGCAGACCGACGAA<br>CGCCGCGGCAACAGGCCGCAGGCGACGGGAGCGGGACGAAGGG | IVT |

**Table S1 (continued)**

| **Name** | **Sequence (5' to 3')** | Source |
|---|---|---|
| IVT template sense promoter | ACGTtaatacgactcactataGG | IDT |
| IVT template CAG 20 scrambled | TGCAattatgctgagtgatatCCCGCTGCGTGCGCGTGTCCGGCGGTGGCCGCTTTCTTTTTGGTCCTGT GTCGTTCCTCGCC | IDT |
| IVT template CAG 31 scrambled | TGCAattatgctgagtgatatCCCTTCGTCCCGCTCCCGTCGCCTGCGGCCTGTTGCCGCGGCGTTCG TCGGTCTGCTTTTCGTTCCTGGGTTGCCTTTGGGTTCGGGCGCGTGTT | IDT |
| IVT template CAG 20 | TGCAattatgctgagtgatatCCGTCGTCGTCGTCGTCGTCGTCGTCGTCGTCGTCGTCGTCGTCGTCG TCGTCGTCGTCGTC | IDT |
| IVT template CAG 31 | TGCAattatgctgagtgatatCCGTCGTCGTCGTCGTCGTCGTCGTCGTCGTCGTCGTCGTCGTCGTCG TCGTCGTCGTCGTCGTCGTCGTCGTCGTCGTCGTCGTCGTCGTCGTC | IDT |
| CAG 31 | CAGCAGCAGCAGCAGCAGCAGCAGCAGCAGCAGCAGCAGCAGCAGCAGCAGCAGCAGCAGC AGCAGCAGCAGCAGCAGCAGCAGCAGCAGCAG | IDT |
| CUG 31 | CUGCUGCUGCUGCUGCUGCUGCUGCUGCUGCUGCUGCUGCUGCUGCUGCUGCUGCUGCUG CUGCUGCUGCUGCUGCUGCUGCUGCUGCUGCUG | IDT |
| CUU 31 | CUUCUUCUUCUUCUUCUUCUUCUUCUUCUUCUUCUUCUUCUUCUUCUUCUUCUUCUUCUUC UUCUUCUUCUUCUUCUUCUUCUUCUUCUUCUU | IDT |
| CAG 3 – A488 | [Alexa 488]-CAGCAGCAG | IDT |
| *Mja*5ext-F | CAAATCAACCTAAAACTTACACAGGCGGGTAAGGGGGCTG | |
| *Mja*3ext-R | CGAACAAAAAGTTTACCATACTCGCGGGTATGGGGGCTATAGC | |
| *Mma*5ext-F | CAAATCAACCTAAAACTTACACATCGTGGCAAGGGGGCTGGTGAC | |
| *Mma*3ext-R | CGAACAAAAAGTTTACCTACTCGAAATGGCATCGGGGCTATAGC | |
| T7ext-F | ATCGCTAATACGACTCACTATAGGCAAATCAACCTAAAACTTACAC | |
| F-ext | CGACGTTGTAAAACGACGGCCAG | |
| R-ext | GGAAACAGCTATGACCATGATTACGCCAAG | |

**Supplementary Table S2.** The partial charges used for atoms in polyphosphate fitted using R.E.D. Server [12].

| **Atoms** | $O2^{T}$ | OS | $O2^{M}$ | $P^{T}$ | $P^{M}$ |
|---|---|---|---|---|---|
| Partial charge | -0.9226 | -0.6235 | -0.9692 | 1.07955 | 1.5619 |

#### Movie legends

**Movie 1.** Phase separation of poly(rU). 1.5 mg/mL poly(rU) with 25 mM Tris-HCl, (pH 7.5 at 25°C) and 500 mM $Mg^{2+}$ underwent thermal cycling via temperature-controlled microscopy. Poly(rU) phase separated reversibly with a UCPT of 25.2 ± 1.2°C (n = 3 replicates).

**Movie 2.** Phase separation of poly(rC). 1.5 mg/mL poly(rC) with 25 mM Tris-HCl (pH 7.5 at 25°C) 100 mM $Mg^{2+}$ underwent thermal cycling via temperature-controlled microscopy. Poly(rC) phase separated reversibly with a LCPT of 57.6 ± 1.9°C (n = 3 replicates).

**Movie 3.** Phase separation of poly(rA). 1.5 mg/mL poly(rA) with 25 mM Tris-HCl (pH 7.5 at 25°C) and 5 mM $Mg^{2+}$ underwent thermal cycling via temperature-controlled microscopy going to 80°C. Poly(rA) phase separated irreversibly with a LCPT of 39.3 ± 8.5°C (n = 3 replicates).

**Movie 4.** Phase separation of poly(rA). 1.5 mg/mL poly(rA) with 25 mM Tris-HCl (pH 7.5 at 25°C) and 5 mM $Mg^{2+}$ underwent thermal cycling via temperature-controlled microscopy going to 34°C. Poly(rA) phase separated irreversibly with a LCPT of 32.7°C in this trial.

**Movie 5.** Thermal cycling of poly(rG). 1.5 mg/mL poly(rG) with 25 mM Tris-HCl (pH 7.5 at 25°C) and 1 mM $Mg^{2+}$ underwent thermal cycling via temperature-controlled microscopy going to 80°C. Poly(rG) remained aggregated at all temperatures.

**Movie 6.** Phase separation of poly(P). 1.5 mg/mL poly(P) with 25 mM Tris-HCl (pH 7.5 at 25°C) and 250 mM $Mg^{2+}$ underwent thermal cycling via temperature-controlled microscopy. Poly(P) phase separated with an LCPT of 35.3 ± 5.0°C (n = 3 replicates).

**Movie 7.** Phase separation of CAGx31 RNA. 10 μM CAGx31 with 25 mM Tris-HCl (pH 7.5 at 25°C), 10 mM $Mg^{2+}$, and 10 mM $Na^{+}$ underwent thermal cycling via temperature-controlled microscopy. Observed LCPT = 66.8 ± 3.9°C (n = 3 replicates).

**Movie 8.** Dynamical arrest via percolation of CAGx31 RNA upon phase separation. 100 μM CAGx31 in a buffer containing 10 mM Tris-HCl (pH 7.5 at 25°C), 50 mM $Mg^{2+}$, and 25 mM $Na^{+}$, underwent phase separation at 38.0 ± 3.5°C (n = 3 replicates) followed by the formation of an extensive percolated network of aspherical droplets. These droplets relaxed and merged into spherical droplets as the temperature was increased to 80°C.

**Movie 9.** Rapid fusion of CAGx31 droplets. 50 μM CAGx31 phase separated at 41.1 ± 1.7°C in a buffer containing 10 mM Tris-HCl (pH 7.5 at 25°C) and 50 mM $Mg^{2+}$. Droplets underwent rapid shape relaxation through coalescence as the temperature was increased past 60°C.

**Movie 10.** Phase separation and arrest of a scrambled CAGx31 sequence. 50 μM scrambled CAGx31 is shown in a buffer containing 10 mM Tris-HCl (pH 7.5 at 25°C), 50 mM $Mg^{2+}$, and 25 mM $Na^{+}$. An irreversible phase transition was observed with a LCPT at 57.9 ± 4.2°C (n = 3 replicates).

**Movie 11.** Phase separation and arrest of CAGx20. 155 μM CAGx20 is prepared in 10 mM Tris-HCl (pH 7.5 at 25°C) and 200 mM $Mg^{2+}$. Irreversible phase separation was observed at 59.9 ± 2.3°C (n = 3 replicates).

**Movie 12.** Reversible phase separation of CUGx31. 50 µM CUGx31 RNA underwent thermal cycling with an LCPT of 72.67 ± 5.4°C in a buffer containing 10 mM Tris-HCl (pH 7.5 at 25°C), 50 mM $Mg^{2+}$, and 25 mM $Na^{+}$. Droplets dissolved after crossing the LCPT as temperature decreased. The same condition for CAGx31 resulted in irreversible droplet formation.

**Movie 13.** Absence of phase separation for CUUx31. 50 µM CUUx31 RNA underwent thermal cycling in a buffer containing 10 mM Tris-HCl (pH 7.5 at 25°C), 50 mM $Mg^{2+}$, and 25 mM $Na^{+}$. No phase separation was observed.

**Movie 14.** Phase separation and percolation of *Pfu* RNase P RNA. 10 µM *Pfu* RPRs were imaged in a buffer containing 50 mM HEPES-KOH (pH 7.5 at 25°C), and 50 mM $Mg^{2+}$. Irreversible phase separation was observed at 68.2 ± 1.3°C (n = 3 replicates).

**Movie 15.** Phase separation and percolation of *Mja* RNase P RNA. 10 µM *Mja* RPRs were imaged in a buffer containing 50 mM Tris-HCl (pH 7.5 at 25°C), and 50 mM $Mg^{2+}$. Irreversible phase separation was observed at 65.9 ± 3.2°C (n = 3 replicates).

**Movie 16.** Phase separation and percolation of *Mma* RNase P RNA. Ten µM *Mma* RPRs were imaged in a buffer containing 50 mM Tris-HCl (pH 7.5 at 25°C), and 50 mM $Mg^{2+}$. Irreversible phase separation was observed at 54.2 ± 0.9°C (n = 3 replicates).

**Movie 17.** Melting of arrested *Pfu* RPR droplets. Droplets were prepared via annealing in a buffer containing 10 mM Tris-HCl (pH 7.5 at 25°C), 25 mM $Mg^{2+}$, and 10 mM $Na^{+}$. Arrested *Pfu* RPR droplets then underwent thermal cycling via temperature-controlled microscopy showing the subsequent relaxation of the *Pfu* RPRs condensates into spherical droplets.

**Movie 18.** FRAP of *Pfu* RPRs. 10 µM *Pfu* with 10 mM Tris-HCl (pH 7.5 at 25°C) and 50 mM $Mg^{2+}$ was used for FRAP experiments.

**Movie 19.** FRAP of *Mja* RPRs. 10 µM *Mja* with 10 mM Tris-HCl (pH 7.5 at 25°C) and 50 mM $Mg^{2+}$ was used for FRAP experiments.

**Movie 20.** FRAP of *Mma* RPRs. 10 µM *Mma* with 10 mM Tris-HCl (pH 7.5 at 25°C) and 50 mM $Mg^{2+}$ was used for FRAP experiments.

**Movie 21.** Reversible phase separation of *Mja* RPRs in a refolding buffer. 20 µM RNA was prepared via a refolding protocol and diluted into refolding buffer containing 50 mM HEPES-KOH (pH 8.0 at 25°C), 10 mM $Mg^{2+}$, and 800 mM AmAc, upon reaching 37°C. Reversible phase separation was observed with an LCPT of 86.3 ± 3.1°C (n = 3 replicates).

**Movie 22.** Reversible phase separation of *Mma* RPRs in a refolding buffer. 20 µM RNA was prepared via a refolding protocol and diluted into refolding buffer containing 50 mM Tris-HCl (pH 7.5 at 25°C), 7.5 mM $Mg^{2+}$, and 500 mM AmAc, upon reaching 37°C. Reversible phase separation was observed with an LCPT of 49.2 ± 3.2°C (n = 3 replicates).

**Movie 23.** Absence of an apparent phase separation of *Pfu* RPRs in a refolding buffer. 20 µM RNA was prepared via a refolding protocol and diluted into refolding buffer containing working concentrations of 50 mM HEPES-KOH (pH 8.4 at 25°C), 10 mM $Mg^{2+}$, and 800 mM AmAc, upon reaching 37°C.

**Movie 24.** Absence of an apparent phase separation of *Pfu* RPRs in a refolding buffer. 2.5 μM RNA was prepared in a buffer containing 50 mM HEPES-KOH (pH 7.5 at 25°C), 10 mM $Mg^{2+}$, and 800 mM AmAc. The sample was heated to 80°C.

**Movie 25.** Dissolution of CAGx31 droplets upon addition of EDTA. 10 μM CAGx31 RNA underwent annealing in a buffer containing 10 mM Tris-HCl (pH 7.5 at 25°C) and 50 mM $Mg^{2+}$. Droplets dissolved upon the addition of a small volume of 500 mM EDTA from the left of the imaging area.

**Movie 26.** Phase separation of CAGx31 RNA in the presence of dimethylsulfoxide (DMSO). Shown here is a solution of 10 μM CAGx31 RNA that underwent thermal cycling in a buffer containing 10 mM Tris-HCl (pH 7.5 at 25°C) and 50 mM $Mg^{2+}$, with the addition of 5% (v/v) DMSO.
